# Expanding the GPCR-RAMP interactome

**DOI:** 10.1101/2023.11.22.568247

**Authors:** Ilana B. Kotliar, Annika Bendes, Leo Dahl, Yuanhuang Chen, Marcus Saarinen, Emilie Ceraudo, Tea Dodig-Crnković, Mathias Uhlén, Per Svenningsson, Jochen M. Schwenk, Thomas P. Sakmar

## Abstract

Receptor activity-modifying proteins (RAMPs) can form complexes with G protein-coupled receptors (GPCRs) and regulate their cellular trafficking and pharmacology. RAMP interactions have been identified for about 50 GPCRs, but only a few GPCR-RAMP complexes have been studied in detail. To elucidate a complete interactome between GPCRs and the three RAMPs, we developed a customized library of 215 Dual Epitope-Tagged (DuET) GPCRs representing all GPCR subfamilies. Using a multiplexed suspension bead array (SBA) assay, we identified 122 GPCRs that showed strong evidence for interaction with at least one RAMP. We screened for native interactions in three cell lines and found 23 GPCRs that formed complexes with RAMPs. Mapping the GPCR-RAMP interactome expands the current system-wide functional characterization of RAMP-interacting GPCRs to inform the design of selective GPCR-targeted therapeutics.

**One-Sentence Summary:** Novel complexes between G protein-coupled receptors and interacting proteins point to a system-wide regulation of GPCR function.

## INTRODUCTION

Protein-protein interactions (PPIs) regulate signal transduction by heterotrimeric guanine-nucleotide-binding regulatory proteins (G proteins). For example, G protein-coupled receptors (GPCRs), which comprise a superfamily of approximately 720 distinct receptors, activate G proteins in response to ligand binding. Receptor activity-modifying proteins (RAMPs) have been shown to regulate GPCR trafficking and ligand specificity for several receptors, including the calcitonin receptor-like receptor (CALCRL) (*1, 2*). Several single-particle cryo-electron microscopy (cryoEM) structures of GPCR-RAMP complexes show them as bimolecular pairs (*1, 3-6*). Bioinformatics studies show that RAMPs globally co-evolved with GPCRs (*7*), and concordance between GPCR and RAMP transcript levels has been observed (*8*). Consequently, elucidation of the GPCR-RAMP interactome has important implications in understanding the cell biology and pharmacology of GPCR signaling and for drug discovery programs that target GPCRs.

We have previously reported an affinity proteomics study using a multiplexed immunoassay based on suspension bead arrays (SBAs). We determined the interactome of 23 GPCRs and the three RAMPs and showed that most secretin family GPCRs interact with at least one RAMP (*9*). In addition, cell-based bioluminescence energy transfer (BRET)-based assay screen have demonstrated that several chemokine GPCRs interact with at least one RAMP (*10*). These studies suggest that GPCR-RAMP interactions might be widespread, but a systematic investigation of the expanded GPCR-RAMP interactome has yet to be reported.

We report the GPCR-RAMP interactome for three RAMPs and 215 GPCRs representing all receptor subfamilies. All potential RAMP-GPCR interacting pairs were expressed ectopically, solubilized and analyzed using the multiplexed SBA strategy. In this assay, large panels of anti-GPCR and anti-RAMP antibodies (Abs) were used to capture and detect GPCR-RAMP complexes on color-coded magnetic microbeads in a flow-based format (*11, 12*). The SBA assay enabled the detection of GPCR-RAMP complexes with up to 11 different capture-detection pairs simultaneously. We found that 122 of the GPCRs tested showed strong evidence for interaction with at least one RAMP. Most RAMP-interacting GPCRs formed complexes with either two or all three RAMPs. However, many GPCRs did not show evidence for complex formation even when co-expressed with a RAMP. We then applied the SBA assay to test for native GPCR-RAMP complexes in three cell lines. We identified 23 GPCRs that formed complexes with at least one RAMP and validated the formation of several native GPCR-RAMP2 complexes by *in situ* proximity assay in neuroepithelioma cells. Most of the specific GPCR-RAMP interacting pairs we report were previously unknown.

## RESULTS

### Workflow to map GPCR-RAMP interactions

We constructed a library of 215 Dual Epitope-Tagged GPCRs (DuET Library) (*12*) and a orthogonal library of three dual epitope-tagged RAMPs. All but four DuET GPCRs have an N-terminal FLAG and a C-terminal 1D4 epitope tag and were derived from the PRESTO-Tango GPCR library (*13*). Each RAMP has an N-terminal 3xHA epitope tag and a C-terminal OLLAS epitope tag. We co-expressed the 215 DuET GPCRs pairwise with each RAMP in Expi293F cells. We coupled anti-epitope tag monoclonal Abs (mAbs), anti-RAMP polyclonal Abs (pAbs), and 248 validated anti-GPCR pAbs primarily from the Human Protein Atlas (HPA) targeting 154 unique GPCRs (*12*) to color-coded magnetic beads. Then, we pooled the beads to create the SBA (**Table S1**). We used six GPCR subfamily-specific SBA pools corresponding to the GRAFS classification system: rhodopsin divided into alpha, beta, gamma, and delta, “other”, and glutamate, secretin, adhesion, and frizzled (GSAF) combined into one group.

We applied the solubilized cell membrane samples containing the co-expressed libraries to the multiplexed SBA (**Fig. 1A**). We detected GPCR-RAMP complexes with either epitope-based or protein-based capture schemes. (**Fig. 1B**). We used five different epitope-based capture schemes: two schemes based on GPCR capture and three based on RAMP capture. In the assay, the GPCR was immunocaptured with anti-1D4 or FLAG mAbs and then the GPCR-RAMP complex was detected via the RAMP using phycoerythrin (PE)-conjugated anti-OLLAS mAb. Similarly, bead-bound Abs captured the RAMP via anti-HA, anti-OLLAS, or anti-RAMP-specific Abs and then the complex was detected via the GPCR using a PE anti-1D4 mAb. Using anti-GPCR pAbs, primarily obtained from the HPA project, the assay captured specific GPCRs and then used PE-conjugated anti-OLLAS mAbs targeting the RAMP to detect the presence of the bead-bound GPCR-RAMP complexes. To detect native GPCR-RAMP interactions in untransfected cell lines, we captured GPCRs with anti-GPCR pAbs and detected the RAMP with PE-conjugated anti-RAMP pAbs. We selected SH-SY5Y cells, SK-N-MC cells, and Expi293F cells for the native screen based on their reported RAMP expression profiles, RNASeq data, and accessibility (*2, 14-16*) [proteinatlas.org].

**Fig. 1.**
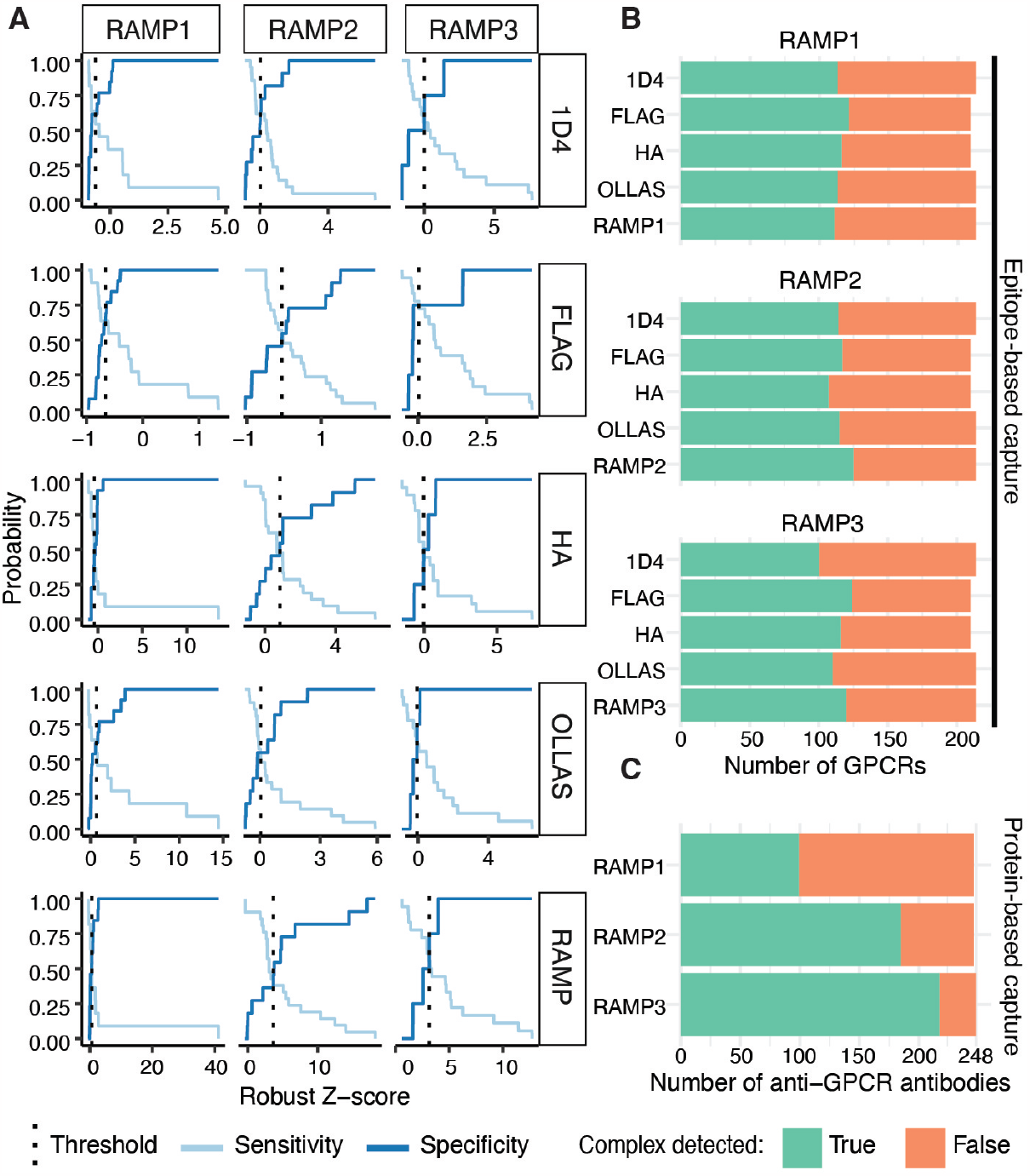
Multiplexed detection of GPCR-RAMP complexes. (**A**) An SBA assay experimental workflow was developed (*9, 12*). Abs were grouped by the phylogenetic subfamily of their GPCR target and were coupled to unique color-coded beads and pooled to generate six subfamily-specific SBAs (1). A library of dual epitope-tagged GPCRs and RAMPs were expressed pairwise. Cells were solubilized to create heterogeneous mixtures of proteins, and concentrations were normalized across samples prior to incubation of aliquots with the SBAs (2). PE-conjugated anti-1D4 or anti-OLLAS mAbs were used to detect the GPCR-RAMP complexes captured by the Ab-coupled beads (3). The data were collected on a Luminex FlexMap 3D instrument (4) and processed to identify GPCR-RAMP complexes. Results were integrated into an interactive web interface (5). (**B**) The GPCR-RAMP complex capture and detection schemes are shown schematically. In all cases, the reporter fluorescence produced by the PE-conjugated detection Ab was associated with the bar code of each bead. From a single well, GPCR-RAMP complexes could be detected simultaneously using either anti-epitope tag Abs, anti-GPCR Abs, or anti-RAMP Abs. (**C**) Data analysis workflow. Data generated as described in (**A**) were first tested for GPCR or RAMP expression, then the fluorescence intensity data were normalized and threshold values were calculated. Adapted from Dahl et al., (*12*). Created in Biorender.com.

To assess the SBA data systematically, we developed a framework that was generalizable to both the heterologous expression and native expression interactome screens and consistent across experiments, yet versatile enough to be customizable to different features of each dataset (**Fig. 1C**). After data collection, we subjected the reported median fluorescence intensity (MFI) levels to several quality control (QC) steps to ensure successful Ab-bead coupling to the beads and to quantify the amounts of solubilized protein added to the assay. We normalized the data where appropriate and transformed it into signal-to-noise ratios (SNRs), Z-scores, or Robust Z-scores (R.Z-scores) for each capture-detection scheme separately. Next, we annotated all potential interactions by setting thresholds for each capture-detection scheme for each RAMP. To enable interactive and user-friendly access to the data, we developed a web-based interface (*Shiny* App, **fig. S1**). The app allows browsing the data in a GPCR-centric manner and includes information about the different data layers reported per GPCR-RAMP interaction. It also summarizes the interactome analysis for each GPCR or ligand subfamily.

### Exploring the GPCR-RAMP interactome

#### Validation of constructs, controls, and SBA assay

To evaluate the suitability of the SBA assay, we employed calcitonin receptor-like receptor (CALCRL) in complex with each of the three RAMPs as positive controls and measured agonist-dependent inositol monophosphate (IP1) accumulation using a homogenous time-resolved fluorescence (HTRF) assay (**fig. S2A, Table S2**). Cells co-expressing the DuET CALCRL (FLAG-CALCRL-1D4) construct and appropriate RAMP were treated with calcitonin gene related peptide (CGRP) or adrenomedullin (AM). The IP1 accumulation responses elicited by these CALCRL-RAMP pairs were compared with cells co-expressing the HA-CALCRL-1D4 construct with each RAMP as used earlier (*9*). We found that the 3xHA- and OLLAS-dual epitope-tagged RAMPs were equally capable of forming functional CALCRL-RAMP complexes compared with the FLAG- and OLLAS-dual epitope-tagged RAMPs used earlier (*9*). All the CALCRL-RAMP1 complexes tested signaled in response to CGRP and AM stimulation, and all the CALCRL-RAMP2 and CALCRL-RAMP3 complexes tested signaled in response to AM stimulation. This confirms prior knowledge about the CALCRL interactome.

Next, we validated the ability of the SBA assay to detect GPCR-RAMP complexes (**fig. S2B**). We generated solubilized cell membrane samples using the DuET CALCRL construct (FLAG-CALCRL-1D4) co-expressed with either 3xHA-RAMP3-OLLAS or FLAG-RAMP3-OLLAS (*9*). We then used the SBA to capture CALCRL-RAMP3 using anti-epitope tag, anti-CALCRL, or anti-RAMP3 Abs. PE-labelled anti-1D4 or anti-OLLAS mAbs were used to detect the captured complexes (**fig. S2B**). The two RAMP3 constructs exhibited similar expression, as determined by anti-RAMP3 specific capture and OLLAS detection. Both constructs showed concordant ability to form CALCRL-RAMP3 complexes when co-expressed with CALCRL. The complex was consistently detected across five of the six capture-detection schemes tested.

To further validate the FLAG-CALCRL-1D4 construct, we generated solubilized membrane samples from cells co-expressing either the DuET CALCRL construct or the HA-CALCRL-1D4 construct used earlier (*9*) with 3xHA-RAMP3-OLLAS (**fig. S2C**). After subjecting these samples to the SBA assay, we saw similar levels of relative CALCRL protein expression and CALCRL-RAMP3 complex formation. Notably 1D4- and OLLAS-based capture performed better than FLAG- and HA-based capture approaches. Taken together, these results confirm the functionality of the dual epitope-tagged RAMP1, RAMP2 and RAMP3 constructs and the robustness of the SBA assay to capture the positive control CALCRL-RAMP3 complex.

To judge the statistical reproducibility of SBA assay measurements, we expressed two GPCRs from different subfamilies with or without a RAMP in biological triplicate. We then performed the SBA assay in technical duplicate (**fig. S3, Table S3**). GPCR class C group 5 member A (GPRC5A) was expressed with or without RAMP2 (**fig. S3A**), and orexin receptor type 2 (HCRTR2) was expressed with or without RAMP3. Four different epitope-based capture-detection strategies were used in parallel to detect the complexes (**fig. S3B**). Anti-HA or anti-OLLAS mAbs were used to capture the RAMP, while PE-conjugated anti-1D4 mAb was used to detect the GPCR in the complex. Conversely, anti-1D4 or anti-FLAG mAbs were used to capture the GPCR, while PE-conjugated anti-OLLAS mAb was used to detect the RAMP in the complex. The one-sided unpaired Wilcoxon test confirmed a statistically significant difference between the MFI levels from the co-expressed GPCR-RAMP and GPCR-mock samples. The GPRC5A-RAMP2 and HCRTR2-RAMP3 complexes have not been reported earlier.

The results described above confirm the suitability and reliability of the multiplexed SBA assay to identify novel GPCR-RAMP complexes in the setting of a validated positive control using both technical and biological replicates. For the DuET library-based GPCR-RAMP interactome screen, we used one biological replicate of each of the four unique GPCR-containing samples (each GPCR alone, and each GPCR with each of the three RAMPs) in two technical replicates. Each technical replicate represented one detection scheme. We analyzed 860 solubilized cell membrane samples along with controls, which corresponded to six 384-well SBA assay plates, to generate approximately 40,000 unique data points.

To determine the expression levels of each RAMP, we used the MFI data arising from capturing the HA tag on the RAMP or capturing the native RAMP sequence and detecting the OLLAS tag (**fig. S4A, Table S3**). We confirmed significantly elevated expression levels for all three RAMPs (p < 0.0001) and observed that each anti-RAMP Ab was specific for its intended RAMP target. A similar strategy was previously used to measure the expression of the GPCRs (*12*). The positive control, CALCRL-RAMP1, was evaluated in more detail. The control complex was analyzed over 10, 11, 20, or 22 technical replicates depending on the capture-detection scheme. It showed highly significant expression levels and complex formation (p < 0.0001) in all combinations compared to mock across all capture-detection schemes (**fig. S4B**,**C**).

#### Detection of GPCR-RAMP complexes using engineered epitope tags

To map the GPCR-RAMP interactome for the 215 receptors tested with the epitope-based capture approach, we computed a threshold for each capture-detection scheme to assign interactions as “hits” for each RAMP (**Fig. 2A, Table S4**) by creating a model based on known interacting and non-interacting pairs in the literature (*1*). We calculated the True Positive (TP), True Negative (TN), False Positive (FP), and False Negative (FN) values at different thresholds to generate sensitivity and specificity curves and determined the threshold at the intersection point of the curves (**Table S4**). The results from applying these thresholds are presented as a binary heatmap for each capture-detection scheme (**fig. S5A**) and can be summed across each column to determine the total number of GPCRs that interact with each RAMP in each capture-detection scheme (**Fig. 2B)**. The number for TP, TN, FP, and FNs for each RAMP in each capture-detection scheme are listed in **Table S5**. Dissimilarities in the thresholds observed between different capture-detection schemes can be explained by experimental differences in binding affinities of the Abs and differences in the published GPCR interactions reported for each RAMP. Overall, there is good agreement across all capture-detection schemes regarding the total number of RAMP-interacting GPCRs identified: 54% of GPCRs tested had measurable complex formation with each RAMP across all five capture-detection schemes. Twenty-three GPCRs formed complexes with all three RAMPs detected by all five capture-detection schemes. Only nine GPCRs did not form any detectable complexes with at least one of the RAMPs in any capture-detection scheme.

**Fig. 2.**
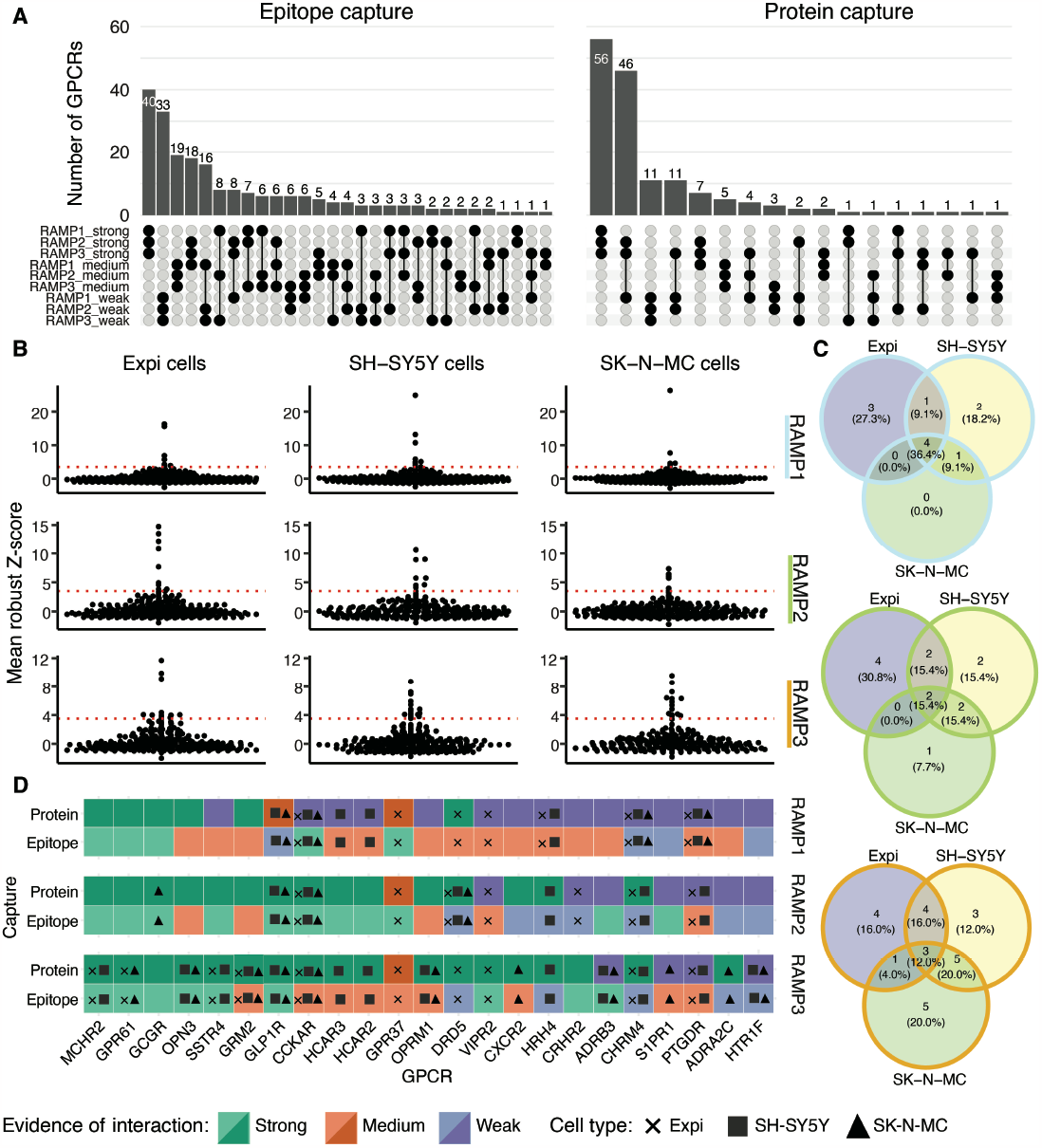
Summary of GPCR-RAMP complex screen. (**A**) Thresholds for each RAMP and epitope-based capture-detection scheme were chosen as the Robust Z-score (R.Z-score) where the sensitivity (light blue) and selectivity (dark blue) curves intersect for GPCR-RAMP interactions known from the literature (*1*). The boxed label on the right of each row indicates the capture scheme. Anti-1D4 and anti-FLAG capture (GPCR capture) corresponds to anti-OLLAS RAMP detection. Anti-HA, anti-OLLAS and anti-RAMP capture corresponds to anti-1D4 GPCR detection. (**B**) The thresholds determined in (**A**) and in **Table S6** were applied to identify GPCR-RAMP interactions. Summary plots of the total number of GPCR-RAMP interactions detected for 215 GPCRs, per RAMP and capture-detection scheme. (**C**) Summary plot of the total number of GPCR-RAMP interactions detected, per RAMP, using protein-based capture with 248 anti-GPCR Abs corresponding to 154 unique GPCRs. Green: complex detected; orange: complex not detected.

#### Detection of GPCR-RAMP complexes using validated anti-GPCR Abs

We used 248 anti-GPCR Abs that recognize 154 unique GPCRs to generate additional evidence for potential GPCR-RAMP interacting pairs (*12*). The protein-based capture approach does not require GPCRs with engineered epitope tags. We selected Ab-specific interaction thresholds (**Table S6**) using a population density-based approach on the R.Z-scores, analogous to that described in the context of the previous Ab validation study (*12*). We selected a strictness of six median absolute deviations (MADs) above the data population density peak. Consistent with the results from epitope-based capture, widespread GPCR-RAMP interactions were detected among the 154 GPCRs tested, with an overall hit rate of approximately two-thirds (**fig. S5B, Table S5**). RAMP1 exhibited the lowest frequency of interactions, where 99 out of 248 Abs (40.0%) captured 74 unique GPCR-RAMP1 complexes. Conversely, we detected 128 GPCR-RAMP2 complexes, corresponding to 185 unique capture Abs (74.6%), and 139 GPCR-RAMP3 complexes, corresponding to 217 unique capture Abs (87.5%).

### Comparison of GPCR-RAMP complex detection schemes

We compared the overall results from epitope-based and protein-based capture formats. First, we assessed whether any capture-detection schemes were subject to bias caused by relative GPCR or RAMP expression levels. Based on the proportion of capture-detection schemes (epitope-based and protein-based capture considered separately) with GPCR-RAMP interactions detected, GPCR-RAMP pairs were classified into three groups of interaction evidence; weak (<33%), medium (>33%, <67%) and strong (>67%). We then examined the GPCR-RAMP interaction “hits” distribution for each GPCR or RAMP expression quartile (**fig. S6, S7**). We did not observe any patterns of interaction evidence correlating with RAMP expression levels (**fig**.**S6A, B, fig. S7A**), indicating that the RAMP expression levels did not bias the complex detection results. We observed fewer high-confidence interactions for RAMP1 in the protein-based capture format and RAMP1 generally expresses less efficiently than RAMP2 or RAMP3. There was a slight tendency of expression bias in the RAMP2 dataset for the protein-based capture format. In contrast, we observed some bias in the distribution of hits across GPCR expression quartiles for the epitope-based but not the protein-based capture formats used to detect GPCR-RAMP complexes (**fig. S6C, D**). Parsing the epitope-based data into individual capture-detection schemes reveals that the staircase distribution can be attributed to capture-detection formats that capture the RAMP (HA, OLLAS, or RAMP specific capture) and detect the GPCR (1D4 tag detection) (**fig. S7B**). The trends seen here indicate that GPCR expression might be limiting in the SBA assay when a tag on the GPCR is used for detection. It is likely that the accessibility of the tag is influenced by immunocaptured complex.

To determine whether RAMP-interacting GPCRs tended to interact with only one RAMP, all three RAMPs, or a subset of the RAMPs, we investigated the overlap between the results for epitope-based and protein-based capture (**Fig. 3A**). For both datasets, the largest group of GPCRs were those with strong evidence for interactions with all three RAMPs – 40 GPCRs in the epitope-based capture set and 56 in the protein-based capture set. There were 20 GPCRs with strong evidence for interaction with all three RAMPs across both datasets. The epitope-based capture results reveal that GPCRs that do not interact with all three RAMPs were next most likely to interact with none of the RAMPs (33 GPCRs). The protein-based capture results show that GPCRs not interacting with all three RAMPs were next most likely to interact with only RAMP2 and RAMP3 (46 GPCRs). We observed this trend earlier (*9*), which could be explained by a lower sensitivity for complex detection via GPCR capture by epitope tag. Notably, interactions unique for single RAMPs were found to be rare.

**Fig. 3.**
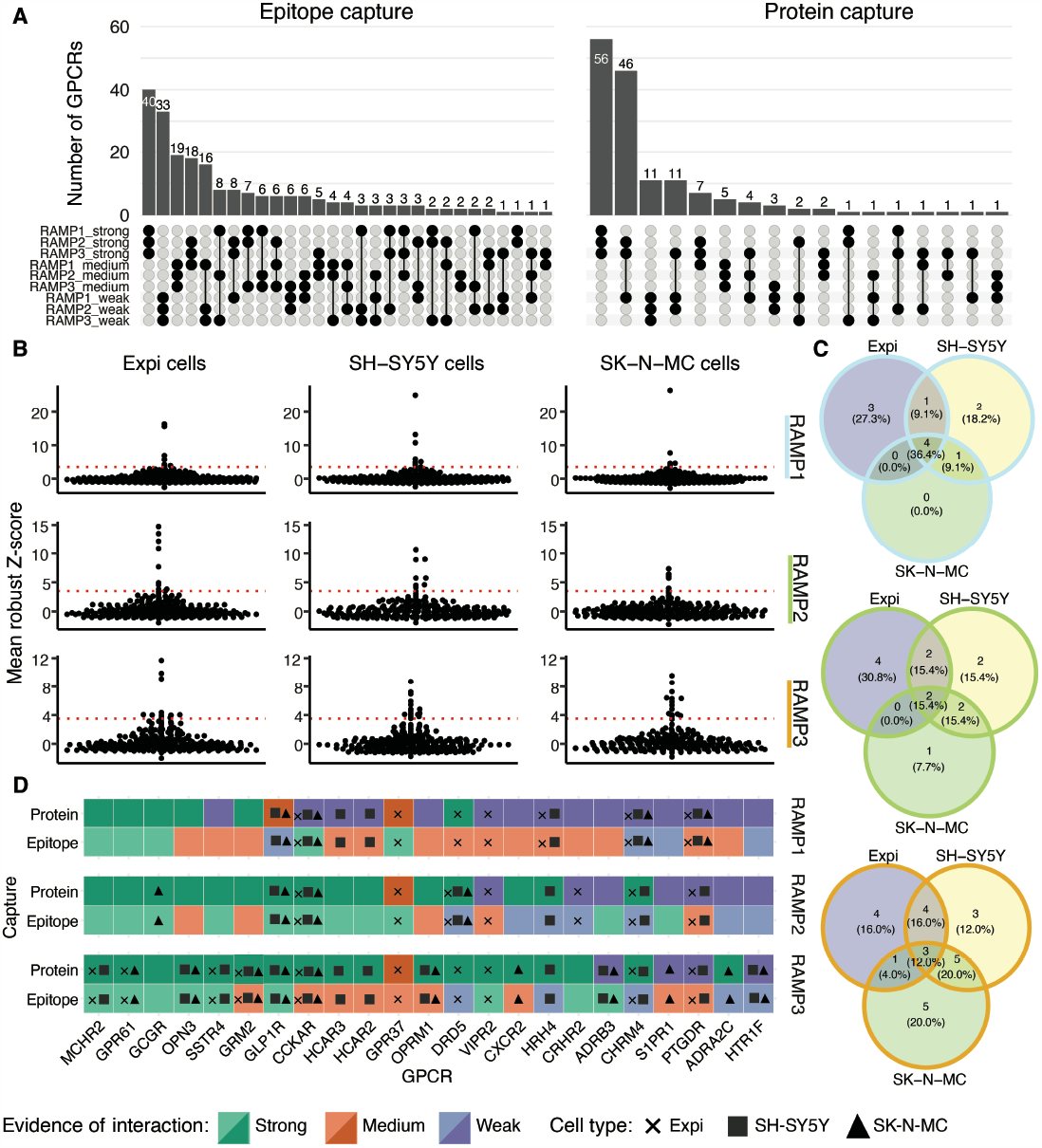
Detection of native GPCR-RAMP complexes in cell lines compared with epitope-tagged complexes in transfected cells. (**A**) UpSet plots showing the overlap between the sets of GPCRs that were found to interact with RAMP1, RAMP2 and RAMP3 for complexes detected with epitope capture (left) or protein capture (right). There were five unique epitope-based capture-detection schemes, and between zero and six unique protein-based capture-detection schemes for each GPCR-RAMP pair studied. Protein capture data were generated for 154 GPCRs (out of 215) for which validated anti-GPCR Abs were available. (**B**) The SBA assay (242 Ab-coupled beads against 148 unique GPCRs) was used to analyze solubilized membranes from Expi293F (Expi), SH-SY5Y and SK-N-MC cells (*12*). Native GPCR-RAMP complexes were detected for each cell line with anti-GPCR capture and either anti-RAMP1 (top), anti-RAMP2 (middle) or anti-RAMP3 (bottom) detection. Data are plotted as the mean R.Z-score measured for each anti-RAMP detection Ab. The GPCR Abs with scores above threshold of 3.5 (dotted red line) are listed in **Table S8**. (**C**) Venn diagrams for anti-GPCR Ab hits across the three cell lines tested for GPCR interaction with RAMP1 (light blue outline), RAMP2 (green outline) and RAMP3 (orange outline). Data are from biological triplicates measured in technical duplicate for each detection scheme. (**D**) Heatmap results for GPCR-RAMP native interactome screen in Expi cells (x), SH-SY5Y cells (filled square), and SK-N-MC cells (filled triangle). Strong: >66% passing capture-detection schemes. Medium: 33-66% passing capture-detection schemes. Weak: <33% passing capture-detection schemes.

Next, we compared the overall results from the multiplexed SBA assay (**Table S7**). Nine GPCRs showed evidence for complex formation with all three RAMPs across all capture-detection methods: GABBR1, GPR143, GPR21, GPR61, HTR4, LPAR2, MTNR1A, OXER1, and P2RY11 (see **Table S1** for corresponding GPCR UniProt IDs). Considering each RAMP individually, 24 GPCRs showed strong evidence for interaction with RAMP1 across all capture-detection methods, 37 GPCRs showed strong evidence for interaction with RAMP2, and 34 GPCRs showed strong evidence for interaction with RAMP3. The intersection of these three sets reveals 58 GPCRs, the number of unique GPCRs with positive interaction evidence in every epitope-based and protein-based capture-detection format for interaction with at least one RAMP.

### Detection of native GPCR-RAMP complexes

We applied the SBA assay to detect natively-occurring GPCR-RAMP interactions in solubilized membranes from Expi293T, SK-N-MC, and SH-SY5Y cell lines. We included Expi293F cells transfected with CALCRL alone or co-transfected with each of the three RAMPs to evaluate the functionality of the assay (**fig. S8, Table S3**). We used two validated anti-CALCRL Abs for capture and PE-conjugated anti-OLLAS mAb, anti-RAMP1 pAb, anti-RAMP2 pAb, and anti-RAMP3 pAb for detection of ectopically-expressed CALCRL-RAMP complexes. As expected, we detected all three CALCRL-RAMP complexes with the CALCRL-OLLAS capture-detection scheme. We detected the correct CALCRL-RAMP complex with CALCRL capture and RAMP-specific detection at high statistical significance compared with the MFI levels from samples derived from cells with CALCRL expressed alone. There were differences in the performances of the schemes used for RAMP detection. A comparison of the SNRs for positive and negative samples within each detection scheme showed that RAMP3 performed better than RAMP2, which in turn performed better than RAMP1. The difference in detection Ab performance may be attributed to different affinities of each anti-RAMP Ab, different relative levels of ectopic expression of each RAMP, or a combination of the two factors.

Next, we validated that we could detect native RAMP expression in each of the three cell lines using anti-RAMP pAbs for both capture and detection (**fig. S9, Table S3**). Three of the five anti-RAMP pAbs used for capture were the same as those used for detection, and two of the anti-RAMP pAbs used for capture were distinct from those used for detection. For the three identical capture-detection anti-RAMP pAbs, each was raised against immunogens of 87-103 amino acids in length. Therefore, we reasoned that the same pAb could be used for capture and detection, as individual Abs typically recognize epitopes of five to seven amino acids (*17*). However, the structural representation of epitopes also affects Ab recognition, so the results must be interpreted carefully (*18*). Two Abs used for capture of RAMP1 or RAMP2 were anti-RAMP pAbs from the HPA. We saw a range of endogenous RAMP expression levels that reached statistical significance compared to the negative control (buffer only). Overall, SK-N-MC and SH-SY5Y cells exhibited higher protein expression levels of a given RAMP than Expi293F cells.

Encouraged by these results, we mapped the native GPCR-RAMP interactome in the three cell lines. We found 11, 13, and 25 unique GPCR-RAMP complexes for RAMP1, RAMP2, and RAMP3, respectively (**Fig. 3B-C, Table S8**). Two anti-cholecystokinin A receptor (CCKAR) Abs captured native CCKAR-RAMP1 and CCKAR-RAMP3 complexes, and two different anti-dopamine receptor D5 (DRD5) Abs captured native DRD5-RAMP2 interactions. Interestingly, in both cases, one of the GPCR-specific Abs captured the GPCR-RAMP complex in all three cell lines, while the second Ab targeting the same GPCR captured the same GPCR-RAMP pair in only one or two of the cell lines. This observation may be explained by different Ab affinity towards the native receptors and different levels of GPCR expression per cell line. Three GPCRs were common among the three cell lines for interactions with RAMP1 or RAMP3, and two GPCRs were shared between all cell lines for interactions with RAMP2 (**Fig. 3D**). Overall, the most significant number of GPCR interactions were identified for RAMP3 and, of those GPCRs, many were also identified as RAMP3-interacting hits in our GPCR-RAMP interactome screen (76%, protein-based capture; 33% epitope-based capture).

### Detection of native GPCR-RAMP complexes in cell membranes

To test for GPCR-RAMP complexes in native cell membranes without heterologous over-expression, we employed a proximity-based assay called the MolBoolean method (*19*). The assay allowed us to quantify selected GPCR-RAMP2 complexes in SK-N-MC cells relative to total GPCR and RAMP2 levels for each unique receptor. The MolBoolean method is based on the proximity ligation assay (PLA) concept. It generates rolling circle amplification products (RCPs) to localize a fluorescence signal *in situ* in a cell membrane environment. However, unlike the PLA, the MolBoolean method enables simultaneous visualization of individual proteins and those forming a complex, and is well suited to provide evidence for the physiological relevance of GPCR-RAMP interactions identified by SBA assay.

We first employed multiple controls to verify that natively-occurring positive control between CALCRL and all three RAMPs could be detected by the MolBoolean assay (**fig S10, Table S3**). Omitting either of the primary Abs before sample processing enabled us to measure primary Ab nonspecific binding. Likewise, omitting all primary Abs enabled us to measure the nonspecific binding of the MolBoolean probes (**fig. S10, Table S3**). The number of RCPs per cell was significantly higher in cells incubated with anti-CALCRL and anti-RAMP primary Abs than in cells that received control treatments. As expected, we observed that many of the puncta corresponding to each RAMP were not in complex with CALCRL, which is consistent with the expectation that RAMPs have many GPCR interacting partners in a typical cell membrane.

We next used the MolBoolean method to test native GPCR-RAMP2 interactions in SK-N-MC cells for eight GPCRs included in the SBA assay screen. We stained SK-N-MC cells with Abs targeting each GPCR or co-stained with an Ab targeting each GPCR and an Ab targeting RAMP2. We stained for CALCRL and RAMP2 as the positive control. Overall, the GPCRs were found to interact with RAMP2 (**Fig. 4, Table S1, S3**). Cells stained for RAMP2 and four of the GPCRs tested exhibited a percentage of overlap RCPs per cell (RCPs corresponding to GPCR-RAMP complexes) that did not differ significantly from the positive control. Although the remaining GPCR-RAMP2 pairs exhibited a statistically significant difference in overlap RCPs per cell compared to CALCRL-RAMP2, complex formation was still observed.

**Fig. 4.**
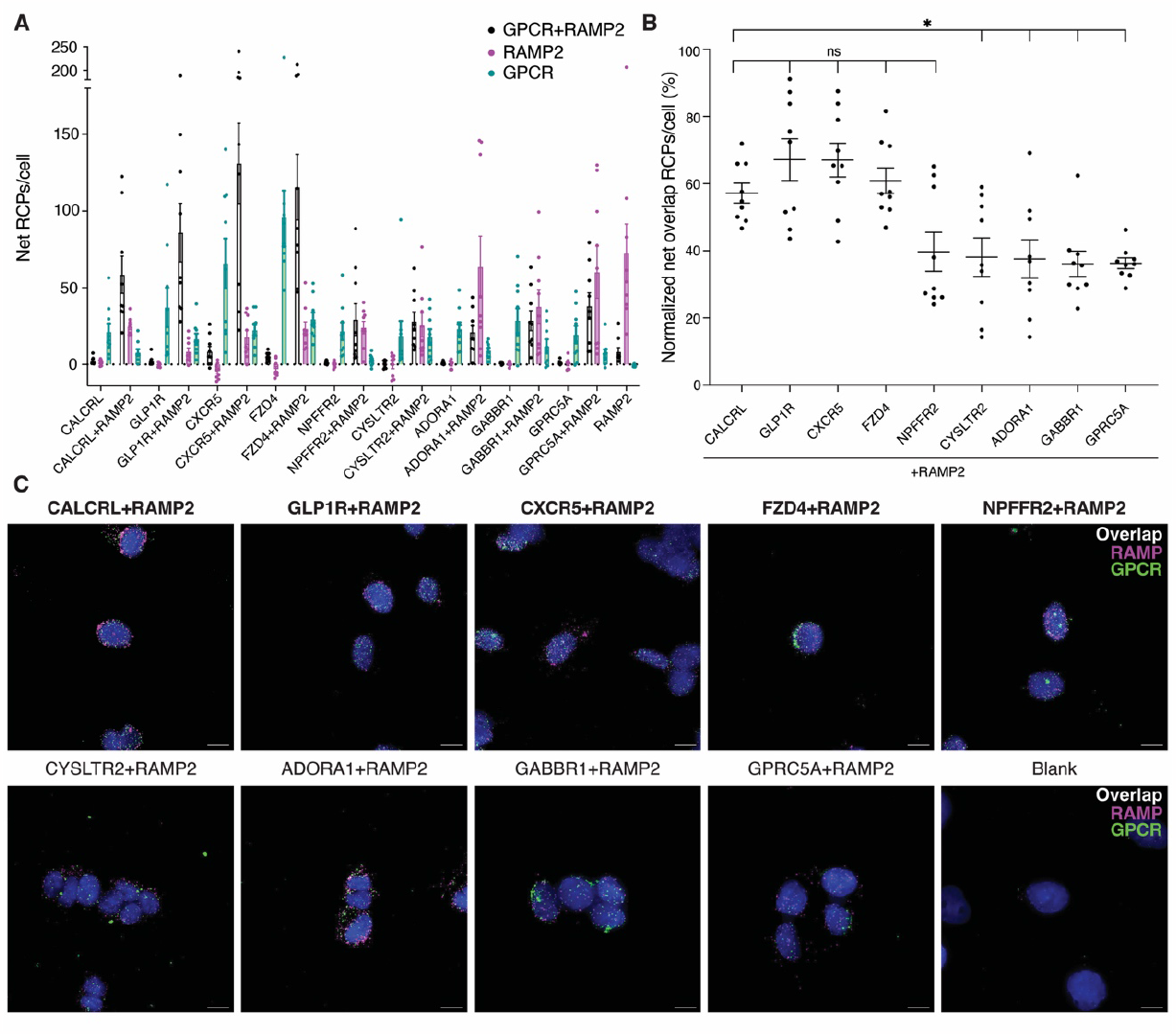
Native GPCR-RAMP2 complexes in SK-N-MC cell membranes. To quantitate GPCR-RAMP2 interactions, the number of MolBoolean rolling circle amplification products (RCPs) per cell for each Z-stack captured from a different field of view was measured. (**A**) The complexed and un-complexed GPCR and RAMP2 are quantified as the number of net RCPs per cell for cells stained for the specified GPCR only, or for the GPCR and RAMP2. (**B**) For each GPCR-RAMP2 pair, the percentage of RCPs out of the total number of RCPs/cell is quantified. Statistical significance was determined by one-way ANOVA followed by Dunnett’s multiple comparisons test to the positive control, CACLRL-RAMP2 (\**p* < 0.05; ns, not significant) Sample sizes and p-values are listed in **Table S3**. (**C**) Representative Z-stack maximum projection images of cells subject to MolBoolean assay. Scale bars, 10 μm. Blue, DAPI; green, GPCR puncta; magenta, RAMP2 puncta; white, GPCR-RAMP2 complex puncta. RCPs from MolBoolean-stained cells that were not incubated with any primary Ab were subtracted to obtain net RCP values. Labels on the x-axis in (**A**), (**B**) and on top of the images in (**C**) indicate the target(s) of the included Abs. The positive control and samples that did not differ significantly from it are bolded. Data are from three biological replicates performed with at least three technical replicates.

## DISCUSSION

We used a combinatorial library and multiplexed screening platform to elucidate the interactome between 215 GPCRs and three RAMPs. The results provide strong experimental evidence to support the hypothesis for widespread GPCR-RAMP interactions across all phylogenetic GPCR subfamilies. Most GPCRs tested interacted with at least one RAMP, and approximately one-quarter of the GPCRs tested interacted with all three RAMPs. Overall, there was satisfactory agreement between the results from epitope-based and protein-based capture of GPCR-RAMP complexes. For 34 anti-GPCR Abs (50 unique GPCR-RAMP pairs, or 7.8% of all potential GPCR-RAMP pairs tested), we observed strong evidence for a particular GPCR-RAMP interaction based on epitope-based capture. However, we failed to observe some of the complexes with protein-based capture. We also investigated the topology corresponding to the antigens for these Abs. We found that extracellular domain (ECD)-transmembrane (TM)1 and extracellular loop (ECL)2-TM5 were the most prevalent features of the epitopes from anti-GPCR Abs. Based on published GPCR-RAMP structures, the RAMPs form contacts with the ECD portion of CALCRL and the calcitonin receptor (CALCR) and TM3, TM4, TM5, and ECL2 of the receptor. Supposing that the solved CALCRL-RAMP and CALCR-RAMP structures are generalizable to other GPCR-RAMP interactions, Abs that would capture the GPCR at any of these topological features may fail to recognize the GPCR if a bound RAMP is present.

The screening of native GPCR-RAMP interactomes in different cell lines is inherently limited by the cell line used because different cells express different GPCRs. Cells typically express at least 100 GPCRs and many of the highest expressed receptors are orphans (*20*). Future work screening tissue samples and cell lines derived from different tissues will likely reveal additional native GPCR-RAMP interactions that we did not detect here. For example, the adenosine A1 receptor (ADORA1) showed strong evidence for RAMP interaction by the SBA assay screen based on ectopic expression, but was not a hit in the native GPCR-RAMP interactome screen. Repeating the screen in a thyroid cancer cell line such as FTC-133, which is also reported to express RAMP1 and RAMP2, may reveal native ADORA1-RAMP interactions that were not present in the cell lines tested (*15*) [proteinatlas.org].

Several GPCR-RAMP interactions detected in the interactome screen were also classified as hits in the native interactome screen. For example, GLP1R-RAMP2 complexes were robustly detected in cell membranes by the MolBoolean proximity assay. The agreement across assays underscores the robustness of our strategy for membrane detergent solubilization. GLP1R has previously been reported to interact with the RAMPs based on studies conducted only in heterologous over-expression systems (*9, 21, 22*). The *in vivo* effects of RAMP1 and RAMP3 double knockdown on the activity of different peptides that target GLP1R and the glucagon receptor (GCGR) have recently been reported (*23*). We show that solubilized, native GLP1R forms complexes with all three RAMPs in SK-N-MC and SH-SY5Y cells and that there are native GLP1R-RAMP2 complexes in membranes in SK-N-MC cells. These results may have important implications for GLP1R pharmacology and the design of therapeutics for metabolic and autoimmune diseases. Moreover, GLP1R has recently been implicated as a multimodal receptor involved in cardiometabolic disease (*24, 25*).

The orphan receptor GPRC5A formed complexes with RAMP2 and RAMP3 that were detected by the SBA assay with strong evidence across both epitope-based and protein-based capture approaches. The presence of GPRC5A-RAMP2 complexes in cell membranes was confirmed by the MolBoolean method. GPRC5A is reported to play a tumor suppressive role, and its downregulation has been implicated in lung, pancreatic, colorectal, and breast cancer pathology, although its endogenous ligand remains unknown (*26-28*). Future investigations of GPRC5A in the presence of RAMP may lead to its successful de-orphanization. Notably, the other three members of the “GPCR family C group 5” subfamily (GPRC5B, GPRC5C, and GPRC5D) also exhibited evidence for complex formation with RAMP2 and RAMP3. Follow-up studies may reveal subtype-specific modes of regulation of the different receptors by the RAMPs.

In summary, combining the DuET GPCR library with multiplexed assays enabled us to elucidate an expanded GPCR-RAMP interactome. The SBA platform was used to detect GPCR-RAMP complexes in solubilized membranes from cells heterologously expressing GPCRs and RAMPs and from cell lines endogenously expressing GPCRs and RAMPs. Several GPCR-RAMP complexes were further investigated in cell membranes *in situ* using a novel proximity assay. Using this multiplexed system and related proximity-ligation tools, we identified at least 50 previously unidentified GPCR-RAMP complexes in cell membranes. Overall, the data strongly suggest the widespread occurrence of GPCR-RAMP complexes among at least one-half of GPCRs tested, supporting a general role for RAMPs in regulating GPCR biology. Our approach is scalable and flexible and can be readily adapted for various basic and translational applications, such as detecting GPCR heterodimers, interactions with regulatory proteins, and screening for pathological anti-GPCR autoantibodies.

## ACKNOWLEDGMENTS

We thank the Human Protein Atlas team and the Affinity Proteomics Unit at SciLife Laboratory for the discussion concerning data analysis. We thank Dr. Thomas Huber for designing some of the genetic constructs used in this study. We thank the Rockefeller University’s Fisher Drug Discovery Resource Center (DDRC) for advice and access to instrumentation (LI-COR Odyssey M (LI-COR), Biotek Synergy Neo, and BioTek Synergy NEO-TRF Hybrid multi-mode reader (BioTek Instruments). We thank Dr. Alison J. North, Dr. Tao Tong, Dr. Priyam Banerjee and Dr. Ved Sharma at the Rockefeller University’s Bio-imaging Resource Center (BIRC) for training, discussion, assistance with the experiments, and access to instrumentation (DeltaVision Image Restoration Inverted Olympus IX-71 microscope and Imaris software). The PRESTO-Tango plasmid kit was a gift from Bryan Roth (Addgene kit # 1000000068).

## AUTHOR CONTRIBUTIONS

Conceptualization: IBK, TPS, JMS

Methodology: IBK, TDC, TPS, JMS, YC

Investigation: IBK, AB, EC, MS, YC

Software: LD

Formal analysis: IBK, AB, LD, YC, EC, TPS, JMS

Visualization: LD, IBK, AB, EC, YC, JMS

Supervision: TPS, JMS, PS

Resources: MU, TPS, JMS

Writing—original draft: IBK, AB, LD, TPS, JMS

Writing—review & editing: IBK, AB, LD, YC, TDC, MU, TPS, JMS, PS, MU

## DATA AND MATERIALS AVAILABILITY

All data needed to evaluate the conclusions in the paper are present in the paper and/or the Supplementary Materials. Other data as well as code used for analysis will be available at a Zenodo repository upon publication in a peer-reviewed journal. Additional visualizations are presented in the *Shiny* app, published online upon journal publication. Abs used in this study are commercially available from different sources through their stated HPA or RRIDs.

## SUPPLEMENTARY MATERIALS

Materials and Methods

Figs. S1 to S10

Legends for Table S1, S3, S6, and S7

Table S2, S4, S5, and S8

References

