## Supplementary Materials for "Expanding the GPCR-RAMP interactome"

**The PDF file includes:**

Materials and Methods
Figs. S1 to S10
Legends for Table S1, S3, S6, and S7
Table S2, S4, S5, and S8
References

### Materials and Methods

Molecular Biology

All constructs were confirmed by sequencing in the forward and reverse directions (T7, BGHR primers) (Genewiz).

#### RAMP constructs

Epitope-tagged, codon-optimized human RAMP1, RAMP2, and RAMP3 constructs were encoded in a pcDNA3.1(+) expression vector. The plasmids encode an N-terminal 3xHA tag (amino acid sequence YPYDVPDYA) following the Signal Peptide (SP, amino acids 1 to 26, 1 to 42, and 1 to 27 for RAMP1, RAMP2, and RAMP3, respectively) and two C-terminal OLLAS tags (amino acid sequence SGFANELGPRLMGK) linked by a flexible spacer that also includes two Strep-tag II peptides (amino acid sequence WSHPQFEKGGGSGGGSGGGSWSHPQFEK). Each 3xHA-RAMP-OLLAS construct was generated using the TagMaster site-directed mutagenesis kit according to the manufacturer's instructions for "long-range mutation" as previously reported (*1*) using the previously validated FLAG-RAMP-OLLAS construct as a starting point (*2*).

#### DuET Library of GPCR constructs for GPCR-RAMP interactome analysis

The DuET library was generated as described in Dahl et al., (*3*). Briefly, epitope-tagged human GPCR DNA constructs were encoded in pcDNA3.1(+). All GPCRs except CALCRL and FZD4, 5, 6, and 10 were generated based on the PRESTO-tango library of signal sequence-FLAG-GPCRs as a starting point. The PRESTO-Tango plasmid kit was a gift from Bryan Roth (Addgene kit # 1000000068). The DuET FLAG-GPCR-1D4 constructs encode the FLAG tag (DYKDDDDA) following the hemagglutinin (HA) SP, MKTIIALSYIFCLVFA. C-terminal components of the original PRESTO-Tango constructs were removed (V2 tail, TEV site, Tta transcription factor) upon the addition of a full-length 1D4 tag (DEASTTVSKTETSQVAPA). The forward primer used for all receptor was CGTTTAAACTTAAGCTTAGCGCCACCATGAAGACGATCATC (5' to 3') and the reverse primer used was TGCTGGCCTCATCGAATTCACCGGTGCGTCCACCGGTATC (5' to 3'). The primers were designed using the NEBuilder Assembly Tool on the NEB website and purchased from Integrated DNA Technologies at the standard desalting grade. The pcDNA 3.1(+) vector encoding the C-terminal 1D4 tag was subject to enzymatic double digestion and purification. FLAG-CALCRL-1D4 with its endogenous SP removed was generated by replacing the HA tag with a FLAG tag in a previously validated FLAG-CALCRL-1D4 construct (*2*). As described earlier, during the construction of the PRESTO-Tango library, 23 GPCRs with an endogenous SP within the coding sequence ended up with a second upstream HA SP in the final plasmid. Therefore, we removed the endogenous SP from 12 GPCRs (F2R, GABBR1, GLP1R, GPR156, GPR37, GPR97, GRM1, GRM2, GRM4, GRM5, GRM6, GRM7), but not from 11 others (CALCR, CD97, CRHR1, CRHR2, GCGR, GIPR, GPR114, GPRC5B, GPRC5C, GPRC6A, VIPR2). There were no FZD receptors in the PRESTO-tango library, so we designed synthetic genes for FZD4, 5, 6, and 10, which were synthesized at Genewiz. The human FZD4, FZD5, FZD6 and FZD10 plasmids encode the 5-HT3a SP in place of the endogenous SP, followed by an HA epitope tag. We introduced a codon for Ala instead the codon for Arg to optimize the Kozak sequence (GCCGCCACCATGG). The C-terminal tail of the engineered receptors was the full-length (18 amino acid) 1D4 mAb epitope tag. (3).

The Gqs5 Gq chimera protein has been described previously (4). All constructs were available in-house.

Cell culture

#### Human embryonic kidney 293T cells

HEK293T cells were cultured in DMEM GlutaMAX supplemented with 10% FBS at 37°C with 5% $CO_2$. Cells were transiently transfected directly 'in plate' with 1 pg of GPCR plasmid per cell unless otherwise specified. The total DNA amount was maintained at 2pg/cell with empty vector pcDNA3.1(+). Briefly, the appropriate amount of plasmid DNA was diluted with FluoroBrite DMEM (Live Cell Fluorescence Imaging Medium, without phenol red). In a separate mixture, 2.5 µL of Lipofectamine 2000 per µg of DNA was diluted in FluoroBrite DMEM and incubated for 5 minutes prior to being combined with the DNA mixture and incubated for 20 minutes. Concurrently, cells were trypsinized, resuspended in 2x supplemented FluoroBrite DMEM

(30mM HEPES, 8mM glutamine, and 20% FBS), and counted. Cells were mixed with the DNA-Lipofectamine 2000-FluoroBrite DMEM mixture and directly plated onto a black, clear-bottom, tissue culture-treated microplate at the cell density of 5,600 cells in 7 µL/well in low volume, LoBase 384-well plates. All microtiter plates were ozone treated and coated with 0.01% poly-D-lysine.

**Expi293F cells**
Expi293F cells, a suspension-adapted cell line derived from HEK293T cells, were cultured and transfected according to the manufacturer's instructions. Briefly, cells were cultured in serum-free Expi293 Medium using culture flasks under constant shaking at 130 rpm at 37°C with 8% $CO_2$. For transfection, cells were counted using a Nexcelom Cellometer Auto T4, diluted to 2,000,000 cells/mL, and allowed to grow overnight. The next day, the cells were counted, diluted to 3,000,000 cells/mL, and 1.25 mL of cells were transferred to each well of a 12-well culture plate. Transient transfections were then performed with the Expifectamine 293 transfection kit (Thermo Fischer Scientific) per manufacturer instructions. Each well of cells was transfected with 4 µL of Expifectamine Reagent and 0.25 µg of GPCR plasmid DNA. Total transfected plasmid DNA was kept constant at 1.5 µg/well with empty vector pcDNA3.1(+). Enhancers were added 18-24 hours after transfection and cells were harvested 72 hours after transfection. Expi293F cells harvested without transfection were grown for 48 hours in 12-well plates (Sarstedt) with an initial density of $3x10^6$ cells/mL in 1.25mL volume.

**SK-N-MC cells**
SK-N-MC cells (ATTC HTB-10, neuroepithelioma cell line) were cultured at 37°C and 5% $CO_2$. SK-N-MCs grown for the SBA assay were maintained in DMEM with GlutaMax and supplemented with 10% FBS, non-essential amino acids, HEPES, penicillin-streptomycin, and sodium pyruvate. Cells were grown in 10 cm round tissue culture dishes (Sarstedt) and harvested at approximately 80% confluency. SK-N-MCs grown for the MolBoolean assay were maintained in EMEM supplemented with 10% FBS and cultured at 37°C and 5% $CO_2$ in a humidified atmosphere until 80-90% confluency, then seeded at a density of 120,000 cells in 0.4 mL per chamber in 8-well chamber slides (NuncLab-Tek) and allowed to grow for 24 hours prior to assay processing.

**SH-SY5Y cells**
SH-SY5Y (ATCC CRL-2266, neuroblastoma cell line) cells were cultured at 37°C and 5% $CO_2$. The cells were maintained in 10% FBS DMEM with GlutaMax and supplemented with non-essential amino acids, HEPES, penicillin-streptomycin and sodium pyruvate. Cells were grown in 10 cm round tissue culture dishes (Sarstedt) and harvested at ~80% confluency.

<u>IP1 accumulation assay</u>

HEK293T cells were transfected with 1 pg/cell of CALCRL DNA, as described above. 24 hours after transfection, IP1 accumulation assay was performed as previously described (*5*). After incubation for two hours, Homogenous Time-resolved Fluorescence (HTRF) reagents and IP1 calibration standards were added and incubated for two hours in the dark at room temperature (RT). Time-resolved fluorescence signals were read on a BioTek Synergy NEO-TRF Hybrid multi-mode reader (BioTek Instruments). Data were collected from three independent experiments performed in technical triplicate.

CALCRL constructs were assayed by co-transfecting HEK293T cells with different tagged versions of CALCRL alone or with each RAMP, and with the promiscuous $G_{qs5}$ chimera protein at a DNA ratio of 1:1:0.5. $G_{qs5}$ is an engineered $G_q$ protein containing the last five amino acid residues of $G_s$, which allows $G_s$-coupled GPCRs to signal through $G_q$ downstream signaling pathways (*4*).

Data reduction, standard calibration, and transformation of HTRF data were performed as previously described (*5*). Normalized IP1 values for CALCRL and RAMP constructs validation assays were calculated relative to the unstimulated mock-transfected cells (set to 0%) and fully stimulated CALCRL (HA-CALCRL-1D4 or FLAG-CALCRL-1D4) co-expressed with FLAG-RAMP3-OLLAS for validation of CALCRL, or HA-CALCRL-OLLAS co-expressed with RAMP3 (FLAG-RAMP3-OLLAS or 3xHA-RAMP3-OLLAS) for validation of each RAMP. These data were fitted to a three parameters sigmoidal dose-response function (Prism 9).

Clarified lysate preparation

Cell membranes were solubilized with *n*-Dodecyl-b-d-maltoside (DM) detergent (Anatrace) to form micelles around membrane proteins and maintain membrane protein structure. Expi293F cells (used for the GPCR-RAMP interactome and the native GPCR-RAMP interactome SBA assays) or SK-N-MC and SH-SY5Y cells (used for the native GPCR-RAMP interactome SBA assay) were harvested and washed twice with cold PBS. Cells were then incubated in solubilization buffer (50 mM HEPES, 1 mM EDTA, 150 mM NaCl, and 5 mM $MgCl_2$ (pH 7.4)) with 1% (w/v) DM and cOmplete mini protease inhibitor (Roche) for two hours at 4°C with nutation. Following solubilization, lysates were clarified by centrifugation at 22,000g for 20 minutes at 4°C. Solubilized lysates were transferred to a microcentrifuge tube, and total protein content was determined by Protein DC assay according to the manufacturer's specifications. Solubilized lysates were flash-frozen prior to storage.

Suspension bead array (SBA) assays

**SBA generation**

Anti-GPCR, anti-RAMP and anti-tag Abs were covalently coupled to color-coded beads (MagPlex, Luminex Corp.), which are magnetic, bar-coded beads defined by a unique combination of infrared and near-red dyes. Each Ab was coupled to a unique bead ID population, as previously described (*6*). In brief, 1.75 µg of each Ab was diluted in MES buffer (100 mM 2-(*N*-morpholino)ethanesulfonic acid (pH 5.0)) to a final volume of 100 µL. The carboxylated surface of the magnetic beads was activated with *N*-hydroxysuccinimide (Pierce) and 1-ethyl-3-(3-dimethylaminopropyl)-carbodiimide (ProteoChem). After 20 minutes of activation, the diluted Abs were conjugated onto the activated carboxylated beads using NHS-EDC chemistry. After one hour of incubation, unbound Abs were washed away using PBS-T 0.05% before adding BRE buffer supplemented with ProClin 300 (Sigma-Aldrich) and overnight incubation. The following day, Ab-coupled beads were subsequently grouped and pooled. The coupling efficiency was determined using anti-rabbit-RPE and anti-mouse-RPE Abs (Jackson ImmunoResearch), as most of the Abs coupled onto beads were rabbit pAbs or mouse mAbs. The data were collected using a FlexMap 3D instrument (Luminex Corp., xPONENT Software, build 4.3.309.1).

**GPCR-RAMP interactome screen**

The SBA assay was carried out in 384-well format, following the same protocol described previously (*3*), which is consistent with our proof-of-concept study (*2*). Clarified protein lysates were diluted to 2 µg/µL in solubilization buffer (described above) with 0.01% (w/v) DM in a 96-well unskirted PCR plate and diluted 3.6 times again in SBA assay buffer such that 12.5 µL of the lysate (25 µg) was combined with 32.5µL of assay buffer to a final volume of 45 µL. The solution was then transferred to a 384-well assay plate containing 5 µL of bead array per well using CyBio SELMA (Analytik Jena).

For the SBA assay to detect native GPCR-RAMP interactions, the following modifications were incorporated into the 384-well assay plate workflow to adapt it to a 96-well format: on the day of the assay, 'wild type' cell lysates were diluted to 4 µg/µL, and any transfected cells were diluted to 2 µg/µL in assay buffer, to maximize signal to noise in the readout.

For all protocols, the lysates and beads were incubated overnight for 16 hours at 4°C. Next, the plates were washed five times with 100 µL (96-well plate assay) or six times with 60 µL (384-well plate assay) of PBS containing 0.05% Tween 20 (PBST) using a BioTek EL406 washer. Next, 50 µL of PE-conjugated anti-tag detection Abs diluted in BRE containing 0.1% DM, 0.1% Tween 20, and 10% rabbit IgG was added per well, and the plate was incubated for one hour at 4°C. The final dilution used for the detection Abs across all experiments were: PE-conjugated anti-1D4, 1:1000; PE-conjugated anti-OLLAS, 1:500. PE-conjugated anti-1D4 and PE-conjugated anti-OLLAS was generated using the anti-1D4 and anti-OLLAS Abs available in-house and a PE-conjugation kit (Abcam). For the SBA assay to determine native GPCR-RAMP interactions, detection was enabled using PE-conjugated anti-RAMP1, anti-RAMP2, and anti-RAMP3 Abs diluted 1:200 or PE-conjugated anti-OLLAS Ab diluted 1:500.

After the incubation, the beads were washed three times with 100 µL (96-well plate assay) or six times with 60 µL (384-well plate assay) of PBST, and then 100 µL or 60 µL of PBST was added to the beads (96-well and 384-well assays, respectively). The fluorescence associated with each bead was measured using a

Luminex FlexMap 3D instrument (Luminex Corp., xPONENT Software, build 4.3.309.1). The data is reported as median fluorescence intensity (MFI).

The number of replicates performed for each SBA assay is as follows: For the GPCR-RAMP interactome screen, one technical replicate was performed for one biological replicate for each unique GPCR-containing sample (i.e., GPCR expressed alone or GPCR co-expressed with RAMP1, RAMP2 or RAMP3). Technical duplicates were included for a subset of samples chosen at random. For the native GPCR-RAMP interactome screen, technical duplicates of biological triplicates for each cell type were performed.

**SBA data analysis**

Data analysis was performed using *R* version 3.6.0 (*7*) with visualization using *ggplot2* (version 3.4.1) (*8*) or base *R* unless stated otherwise. Expression of transfected RAMP was evaluated by comparing samples with mock transfection with samples from RAMP1-, RAMP2- or RAMP3-transfected cells. The comparisons were made using one-way ANOVA (aov function of the stats package, version 3.6.0) followed by Dunnett's multiple comparison tests (DunnettTest function of the DescTools package, version 0.99.43) separately for capture using anti-HA, anti-RAMP1, anti-RAMP2, and anti-RAMP3 Abs.

To validate the positive control complex CALCRL-RAMP1, a one-way ANOVA followed by Dunnett's multiple comparison tests was performed, comparing mock-transfected cells and CALCRL-RAMP1-transfected cells with empty samples (buffer only). CALCRL expression was validated with anti-FLAG capture and PE anti-1D4 detection. RAMP1 expression was validated with anti-HA capture and anti-OLLAS detection. Complex formation was determined by examining data from anti-RAMP1, anti-HA, or anti-OLLAS capture and anti-1D4 detection or anti-CALCRL, anti-FLAG, or anti-1D4 capture and anti-OLLAS detection.

Reproducibility in detecting GPCR-RAMP complexes was tested by comparing anti-OLLAS or anti-HA capture measurements with anti-1D4 detection or anti-1D4 or anti-FLAG capture with anti-OLLAS detection. The samples tested were solubilized membranes of cells expressing GPCR only or co-expressing a GPCR with RAMP2 or RAMP3. GPRC5A was tested with RAMP2, and HCRTR2 was tested with RAMP3 in biological triplicates and technical duplicates. Differences between GPCR alone and GPCR-RAMP samples were evaluated using the Wilcoxon rank-sum test (wilcox.test function of the stats package) per GPCR, per capture-detection scheme.

To put measurements from different Abs on a similar scale, raw MFI values were transformed to R.Z-scores separately per capture-detection scheme with the formula (x - median(x) / (1.4826 * MAD(x)), with x being the measurement values from one Ab and MAD the median absolute deviation.

GPCR-RAMP complex detection with the five different epitope-based capture-detection schemes was determined by applying a threshold for complex detection, which was defined by constructing specificity-selectivity plots. We considered RAMP-specific capture with anti-RAMP1, RAMP2, and RAMP3 pAbs an "epitope-based" scheme as it was generalizable to all GPCRs tested. The specificity and selectivity curves were plotted as a function of the threshold and displayed as R.Z-score. Sensitivity is defined as the fraction of known interacting GPCRs that were correctly detected as interacting, and specificity is defined as the fraction of known non-interacting GPCRs that were correctly detected as not interacting, using all previously published reports on GPCR-RAMP interactions (*9*). The intersection of the two curves was assigned as the threshold. Detected interactions per capture-detection scheme and GPCR were visualized in a heatmap generated using the *ComplexHeatmap* package (version 2.2.0) (*10*).

GPCR-RAMP complex detection with protein-based capture-detection schemes was analyzed by individually defining a passing threshold for complex detection for each anti-GPCR Ab. The threshold was 6 MADs (constant of 1.4826) of the expected negative population for each anti-GPCR Ab added to the population density peak (*3, 11*). Results were visualized in a heatmap using the *ComplexHeatmap* package (*10*).

To put epitope-based capture and protein-based capture data on the same scale for comparisons, GPCR-RAMP interactions were classified by the fraction of epitope-based capture schemes or protein-based capture schemes (capture with anti-GPCR Abs) that detected an interaction. An interaction was classified as strong if more than 2/3 of the capture-detection schemes passed, medium if the fraction of passing capture-detection schemes was between 1/3 and 2/3, and weak if less than 1/3 of the capture-detection schemes passed.

To investigate whether there were any biases in the detected GPCR-RAMP interactions arising from differences in GPCR or RAMP expression, the proportions of interaction evidence classes (strong, medium, weak) were visualized across quartiles of GPCR or RAMP expression in bar plots. The visualization was also made for epitope capture schemes separately for proportions of passing or failing interactions.

The amount of strong, medium or weak GPCR interactions per was visualized in UpSet plots (ComplexUpset package, version 1.3.3) (12, 13), separately for epitope-based and protein-based capture (10).

**Data analysis of native GPCR-RAMP interactomes**

The MFI data were transformed to signal-to-noise ratios (SNRs) by dividing signals per capture and detection Ab by the median of the corresponding buffer signals. Native interactions were evaluated per cell type and detection Ab. SNRs were normalized using quantile normalization (*normalizeBetweenArrays* function of the *limma* R package, version 3.42.2) to account for potential differences in protein concentration between replicates and then transformed to R.Z-scores. Any data points above 3.5 (*14, 15*) were annotated as interactions.

<u>MolBoolean Assay</u>

**Assay protocol**

SK-N-MC cells seeded onto chamber slides were fixed with 3.7% FA and permeabilized with 0.2% (v/v) TritonX-100 after 24 hours, as described in Raykova et al. (*16*). After fixation, cells were washed twice in TBS and then processed following the instructions for the MolBoolean assay kit provided by Atlas Antibodies (protype product based on Raykova et al., gift from Atlas Antibodies). Although the original MolBoolean assay was designed for mouse and rabbit primary Abs, we used a MolBoolean kit developed by Atlas Antibodies for use with sheep and rabbit primary Abs. The Abs for staining endogenous GPCRs and RAMPs (**Table S1)** were used at a final concentration of 0.002 mg/mL for GPCR Abs and 0.015mg/mL for RAMP Abs. Samples were incubated with primary Abs overnight at 4°C. After MolBoolean processing, cells were mounted in DuoLink mounting medium with DAPI (Sigma-Aldrich), incubated at RT, stored overnight at −20°C, and imaged the following day.

**Image acquisition and processing**

Deconvoluted images were acquired with a DeltaVision Image Restoration Inverted Olympus IX-71 microscope on the blue, red, and far-red channels using a 60x oil immersion objective. Excitation/emission wavelengths are 390 ± 18/435 ± 48 nm for the blue channel (DAPI), 575 ± 25/632 ± 60 nm for the red channel, and 632 ± 22/ 676 ± 34 for the far-red channel. Exposure times and transmittance percentages were constant while imaging all samples in the same experiment. At least three Z-stack images (0.2 μm thickness per slice) of different fields of view were captured per coverslip for each sample.

Image processing was done in ImageJ (adding scale bars, generating maximum projections, and generating split channel images) and CellProfiler (17). Quantification of rolling circle amplification products (RCPs) per channel and quantification of overlapping RCPs was carried out in CellProfiler with a pipeline developed by Raykova et al. and provided by Atlas Antibodies (16, 17). The same pipeline parameters were used for all samples within an experiment (threshold = 0.5 for the RAMP staining and 0.25-1.0 for the GPCR staining). Nuclei were quantified manually. Adjustments for brightness and contrast in ImageJ were made on images for visualization purposes only.

**MolBoolean data analysis**

Total RCPs for each Z-stack were divided by the total number of cells per image, and the number of RCPs from the “blank” sample (cells that were not stained with any primary Ab) was subtracted to obtain net RCPs/cell (Excel). To calculate net normalized RCPs/cells, the number of RCPs corresponding to complexed and un-complexed proteins was normalized to the total number of RCPs/cell. Data with a negative value for net total RCPs/cell were excluded. The net RCPs/cell and net normalized RCPs/cell data were plotted in Prism 9 (GraphPad). Statistical significance was determined by a one-way ANOVA followed by Dunnett’s multiple comparisons test to the positive control, CALCRL-RAMP2 using *R* version 3.6.0 (*aov* function of the *stats* package, version 3.6.0 and *DunnettTest* function of the *DescTools* package, version 0.99.43) (*7*). Outliers were determined in Prism via the ROUT method with $Q$ = 1%. Two outliers were removed from the CALCRL-RAMP2 dataset and one from the CYSLTR2 dataset (net RCPs/cell). Two outliers were removed from the GABBR1-RAMP2 dataset and one from the GPRC5A-RAMP2 dataset (% normalized net overlap RCPs/cell).

Interactive web interface

A web-based companion R app was made using the *shiny* package (version 1.7.1) and R version 4.2.0. The *Shiny* app was created to contain information on interactions for individual GPCRs, which were visualized in heatmaps (*ComplexHeatmap* package, 2.14.0), circle plots (*circlize* package, 0.4.15) (*18*), and density plots (*ggplot2* package, 3.3.6) (*8*), Summaries of interactions were visualized in heatmaps (*plotly* package, 4.10.0) (*19*). The interface was containerized with *Docker* (version 20.10.16, build aa7e414) and hosted online on the *shinyapps.io* platform.

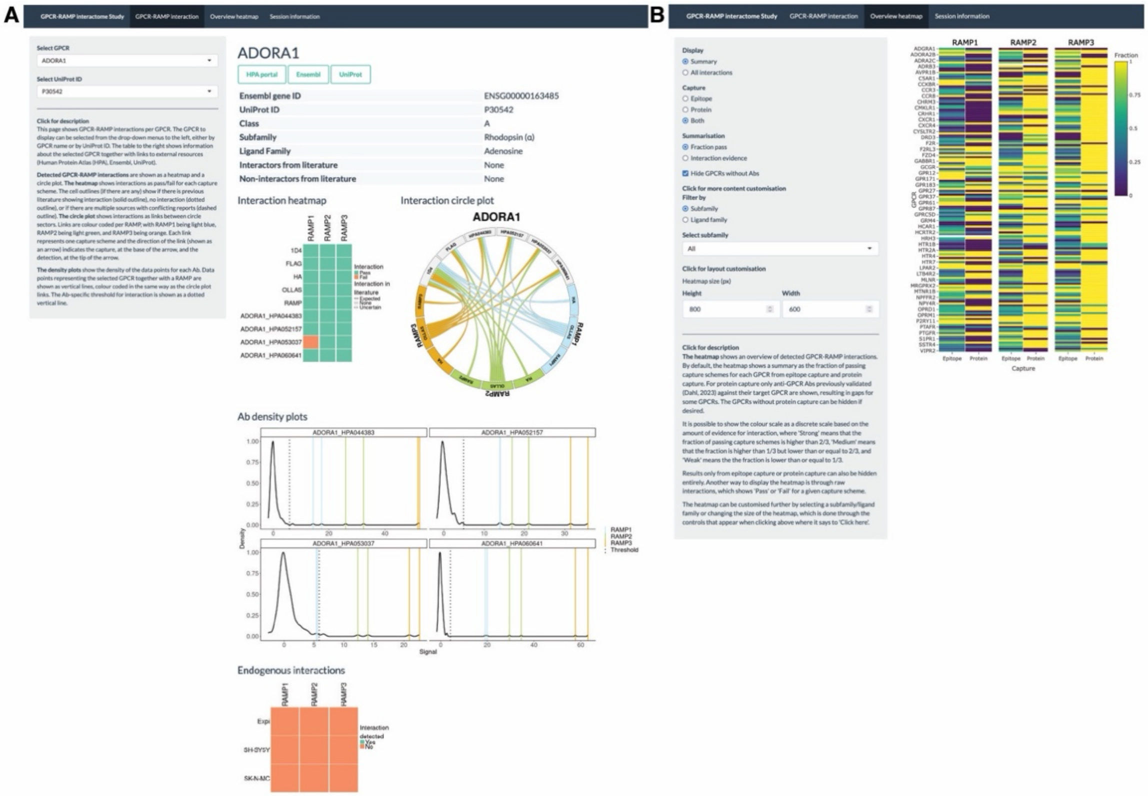


**Fig. S1. Web interface to browse the GPCR-RAMP interactome.**

The results from the GPCR-RAMP interactome analysis are available in an R-based Shiny app that can be accessed in a web browser. (**A**) As exemplified by the entry for the adenosine A1 receptor (ADORA1), the interface shows a summary of GPCR-RAMP interactions detected for all possible capture-detection schemes. The results are displayed as a binary heatmap and alluvial plot, and the density plots for the GPCR-specific capture of GPCR-RAMP complexes with anti-GPCR Human Protein Atlas (HPA) Abs are shown below. The results from the native GPCR-RAMP interactome SBA assay are displayed as a binary heatmap at the bottom. A similar display is available on the app for each GPCR tested. (**B**) The app allows the user to customize the analysis of the GPCR-RAMP interactome dataset to create heatmap displays.

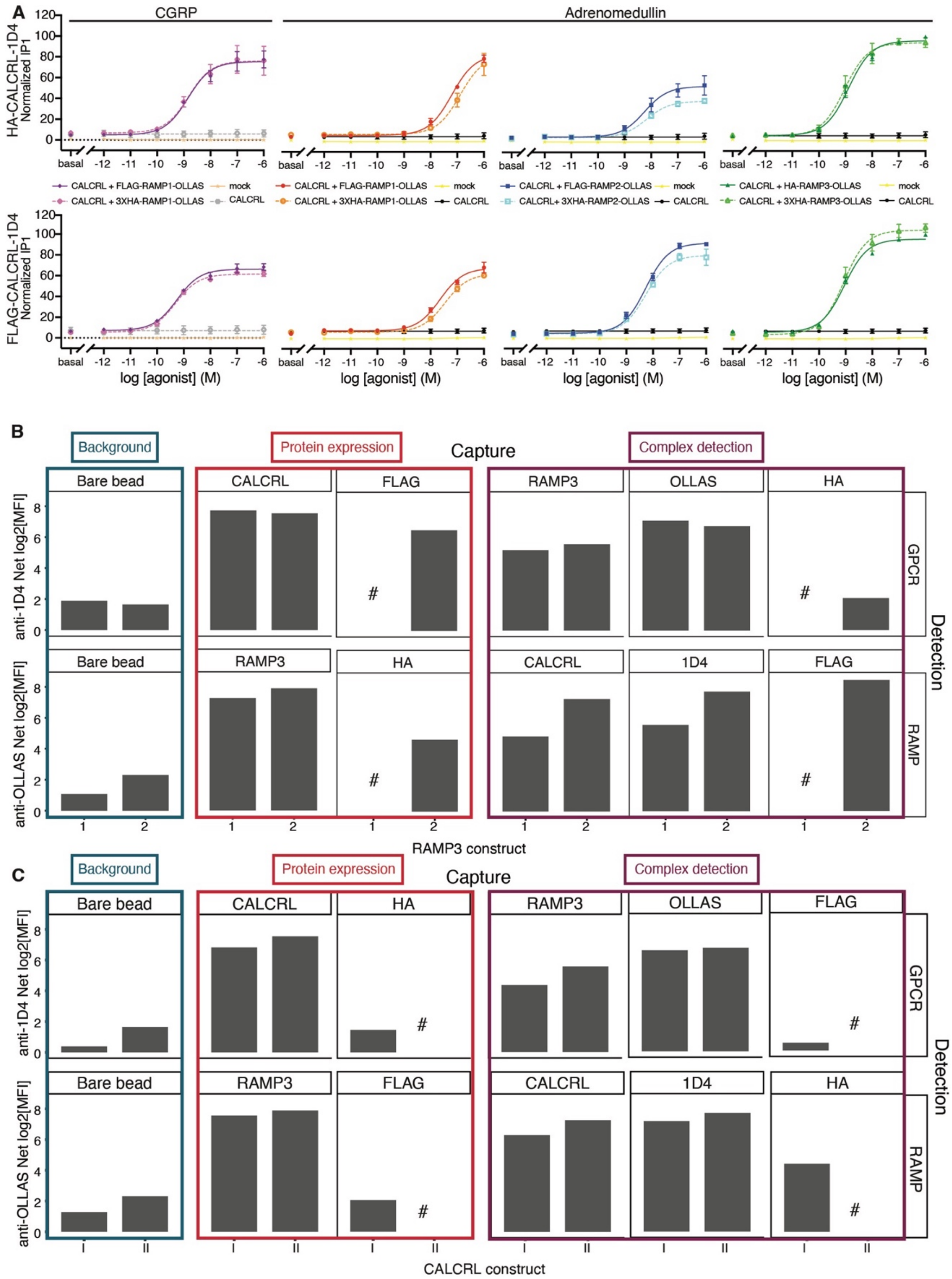
A
CGRP
Adrenomedullin
HA-CALCRL-1D4 Normalized IP1
FLAG-CALCRL-1D4 Normalized IP1
basal
log [agonist] (M)
CALCRL + FLAG-RAMP1-OLLAS
CALCRL + 3XHA-RAMP1-OLLAS
mock
CALCRL
CALCRL + FLAG-RAMP2-OLLAS
CALCRL+ 3XHA-RAMP2-OLLAS
CALCRL + HA-RAMP3-OLLAS
CALCRL + 3XHA-RAMP3-OLLAS
B
Background
Protein expression
Capture
Complex detection
Bare bead
CALCRL
FLAG
RAMP3
OLLAS
HA
1D4
anti-1D4 Net log2[MFI]
anti-OLLAS Net log2[MFI]
GPCR
RAMP
Detection
RAMP3 construct
C
CALCRL construct

**Fig. S2. Validation of GPCR-RAMP complexes used as positive controls.**

(**A**) IP1 accumulation assays were carried out in HEK293T cells expressing each RAMP construct used in this study (3xHA-RAMP-OLLAS) or each of the previously validated RAMP constructs (FLAG-RAMP-OLLAS) (*2*) in the presence of dual epitope-tagged CALCRL (top row, HA-CALCRL-1D4 or bottom row, FLAG-CALCRL-1D4). Dose-response curves were determined for the specified CALCRL with adrenomedullin (all RAMPs) or calcitonin gene-related peptide (CGRP) (RAMP1 only). IP1 accumulation is normalized to maximal adrenomedullin stimulated- HA-CALCRL-1D4 + RAMP3 (top row) or FLAG-CALCRL-1D4 + RAMP3 (bottom row). Data are expressed as the mean ± SEM of normalized IP1 accumulation from three independent experiments performed in four technical replicates. Fitting parameters are provided in **Table S2**. (**B, C**) Constructs described in panel **A** perform similarly in SBA assay format. Expi293F cells were co-transfected with FLAG-CALCRL-1D4 and either FLAG-RAMP3-OLLAS (Construct 1) (*2*) or 3xHA-RAMP3-OLLAS (Construct 2) (**B**) or co-transfected with either HA-CALCRL-1D4 (Construct **I**) (*2*) or FLAG-CALCRL-1D4 (Construct **II**) and 3xHA-RAMP3-OLLAS (**C**). Cells were then solubilized and incubated with the SBA, which included beads conjugated to each of four mAbs targeting HA, 1D4, FLAG, and OLLAS and two previously validated pAbs targeting CALCRL and RAMP3 (*2*). CALCRL-RAMP3 complexes were captured onto the beads, and either CALCRL, RAMP3 or the CALCRL-RAMP3 complex was detected by PE-conjugated anti-1D4 mAb (top row) or anti-OLLAS mAb (bottom row). Dark turquoise: background level (bare bead capture, 1D4, and OLLAS detection). Red: protein expression (CALCRL expression: CALCRL capture, 1D4 detection and FLAG capture, 1D4 detection. RAMP3 expression: RAMP3 capture, OLLAS detection, and HA capture, OLLAS detection). Plum: CALCRL-RAMP3 complex detection (RAMP3 capture, 1D4 detection; OLLAS capture, 1D4 detection; HA capture, 1D4 detection; CALCRL capture, OLLAS detection; 1D4 capture, OLLAS detection; and FLAG capture, OLLAS detection). # indicates that the capture-detection scheme data are omitted for clarity since the tag being captured (FLAG or HA) is present on either CALCRL, and RAMP3, or neither. The y-axis represents the 2-fold change in median fluorescence intensity (MFI) compared to samples with mock-transfected lysate.

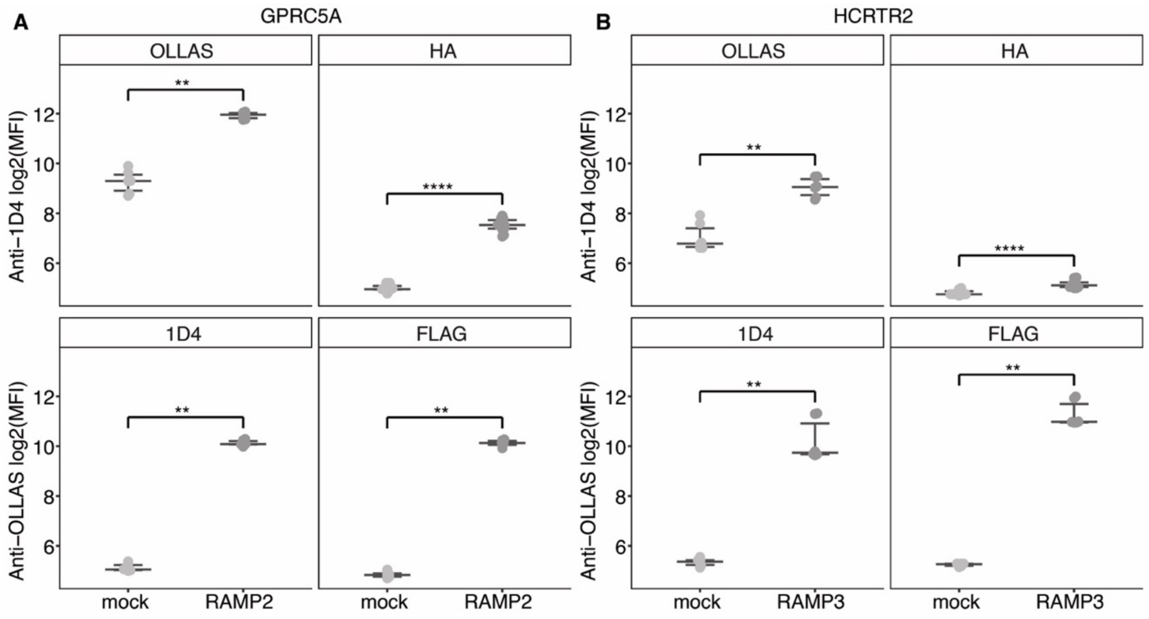


**Fig. S3. Reproducibility of the SBA method to detect GPCR-RAMP complexes.**

Representative examples of GPCRs are shown, where we tested the reproducibility of the SBA assay for detecting solubilized GPCR-RAMP complexes. GPCRs selected from two subfamilies were expressed with or without a RAMP in biological triplicate, and each sample was quantified in technical duplicate (N=6). GPCR class C group 5 member A (GPRC5A; glutamate subfamily) was expressed with or without RAMP2 (**A**), and orexin receptor type 2 (HCRTR2; beta subfamily) was expressed with or without RAMP3 (**B**). Each complex was detected using four epitope-based capture-detection strategies in parallel. *Top row*: the RAMP was captured with beads coupled to anti-HA or anti-OLLAS mAb, and the GPCR was detected with PE-conjugated anti-1D4 mAb. *Bottom row*: the GPCR was captured with beads coupled to anti-1D4 or anti-FLAG mAb, and the RAMP was detected with PE-conjugated anti-OLLAS mAb. Significance was determined by a one-sided unpaired Wilcoxon test (**** $p < 0.0001$, ** $p < 0.01$). Sample sizes and p-values are listed in **Table S3**. X-axis labeling: mock indicates that only the GPCR was ectopically expressed; RAMP2 (**A**) or RAMP3 (**B**) indicate that the GPCR was co-expressed with the specified RAMP. Data are plotted as the log2 of median fluorescence intensity (MFI), and horizontal bars represent the medians of each group with the 25$^{th}$ and 75$^{th}$ percentiles.

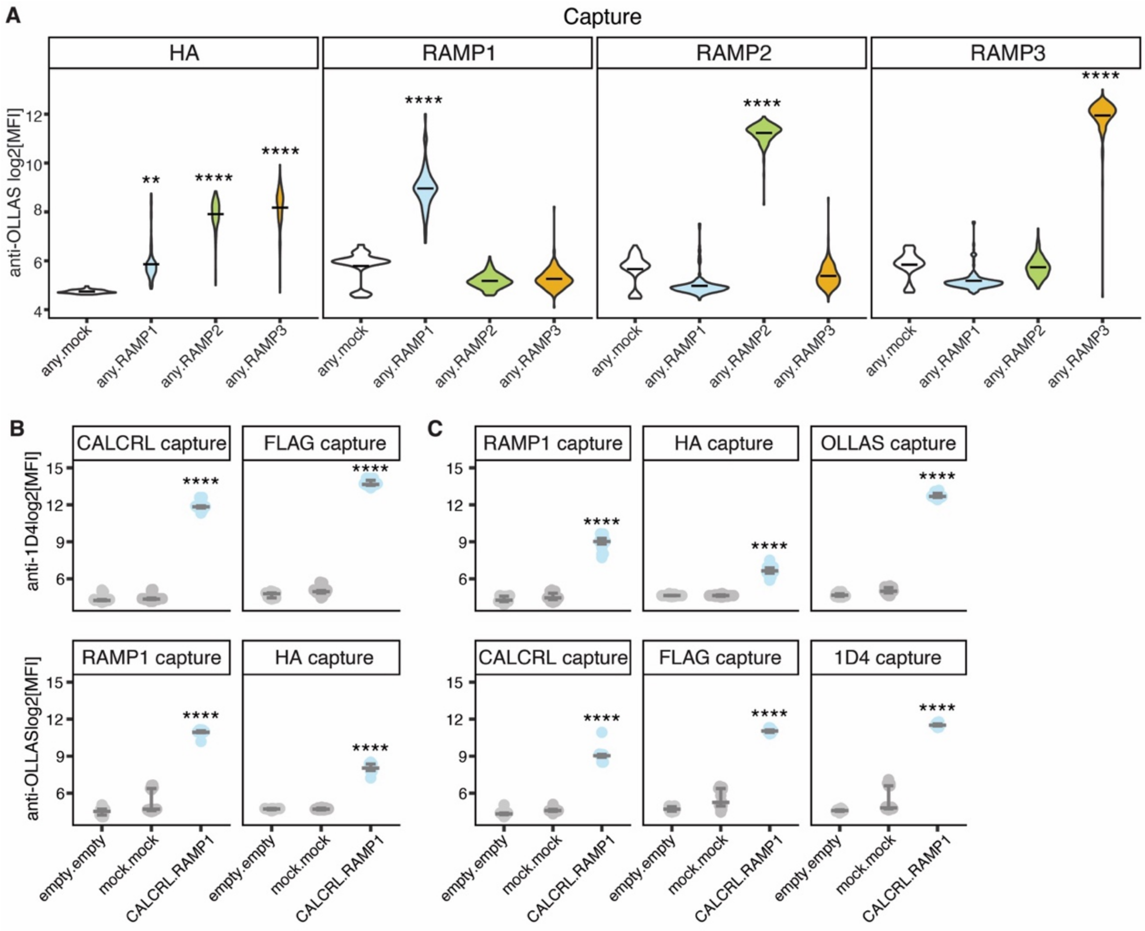


**Fig. S4. RAMP expression and CALCRL-RAMP1 complex formation.**

(**A**) Quantification of expressed RAMPs. Relative quantities of solubilized dual epitope-tagged RAMP 1, 2 and 3 were determined by SBA assay. Each RAMP was captured with beads coupled to anti-HA mAb or anti-RAMP specific pAb capture target as noted at the top of each plot and detected with PE-conjugated anti-OLLAS mAb. Violin plot horizontal lines indicate the mean. Significance was determined by a one-way ANOVA (with $p < 0.05$) followed by Dunnett's multiple comparison test to any.mock (**** $p < 0.0001$, ** $p < 0.01$, if not marked $p >= 0.05$). White, any.mock; light blue, any.RAMP1; lime, any.RAMP2; orange, any.RAMP3. (**B, C**) Validation of expression (**B**) and complex detection (**C**) of the positive control used for the SBA assay screen, CALCRL-RAMP1. Lysates from Expi293F cells co-transfected with epitope-tagged CALCRL and epitope-tagged RAMP1 were incubated with the SBA and CALCRL-RAMP1 complexes were captured on the beads in a multiplexed fashion. The proteins were detected by PE-conjugated anti-1D4 mAb (top row) or PE-conjugated anti-OLLAS mAb (bottom row). (**B**) CALCRL expression (top row) and RAMP1 expression (bottom row) were measured with the capture-detection schemes shown. (**C**) CALCRL-RAMP1 complex formation was measured with six capture-detection schemes: RAMP1 was captured with beads coupled to anti-RAMP1 pAb, anti-HA mAb or anti-OLLAS mAb, and the CALCRL-RAMP1 complex was detected with PE-conjugated anti-1D4 mAb. CALCRL was captured with beads coupled to anti-CALCRL pAb, anti-FLAG mAb or anti-1D4 mAb, and the CALCRL-RAMP1 complex was detected with PE-conjugated anti-OLLAS mAb. Statistical significance was determined by ordinary one-way ANOVA followed by Dunnett's multiple comparisons test to mock.mock (**** *p* < 0.0001, if not marked $p >= 0.05$). The thick horizontal lines represent the median values, and the thin lines above and below show the 75th and 25th percentiles, respectively. Sample names on the x-axis use the format "transfected GPCR name (if any).transfected RAMP name (if any)". Data are plotted as the log2 of median fluorescence intensity (MFI). Sample sizes and p-values are listed in **Table S3**.

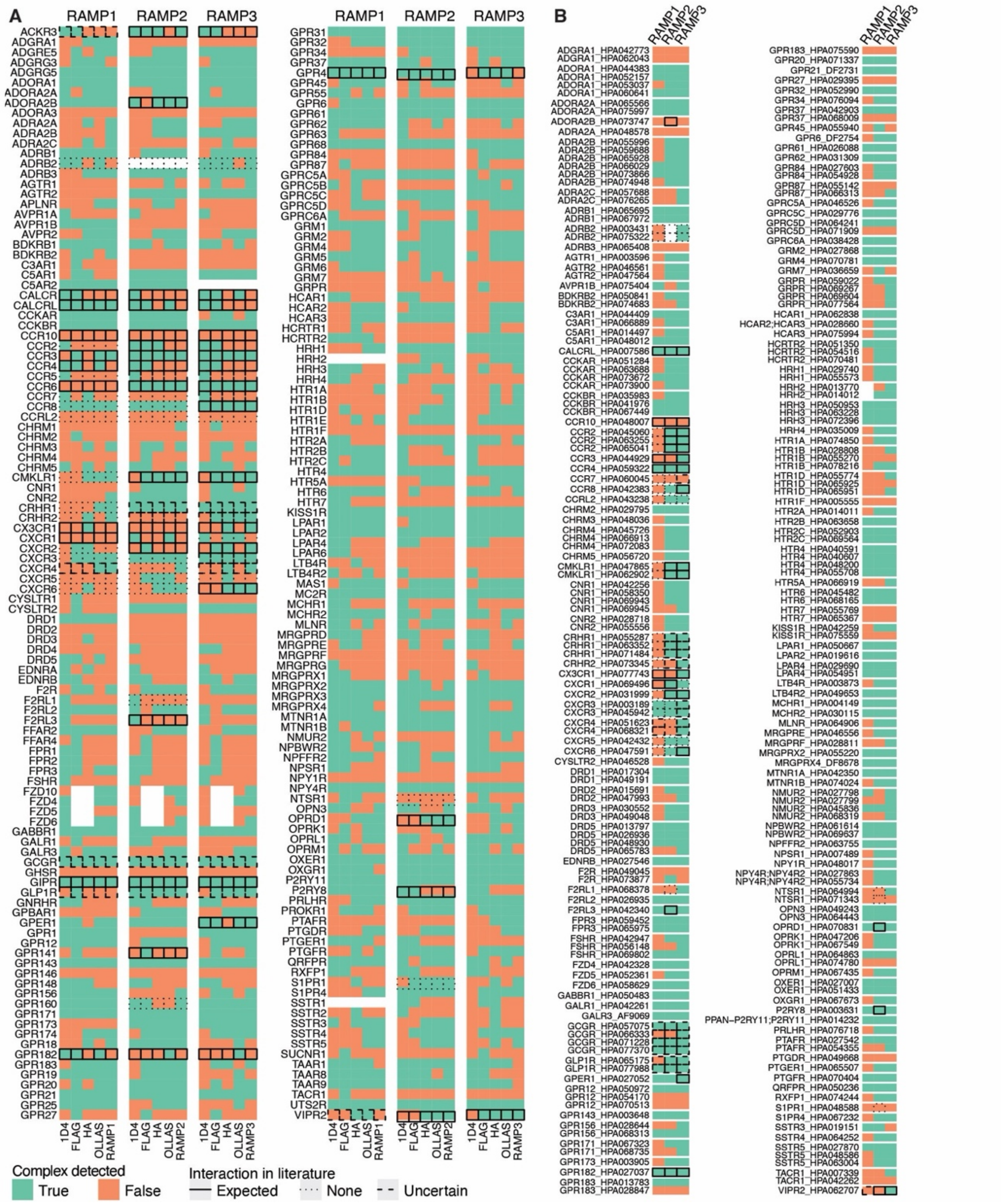


**Fig. S5. Summary of GPCR-RAMP complex data.**

Heatmaps are shown for GPCR-RAMP interactions identified for each RAMP using multiple capture-detection schemes. (**A**) Epitope-based capture. Each row represents a single GPCR. (**B**) Protein-based capture. Each row represents a single anti-GPCR Ab labeled with the format "GPCR target_Ab code". Different Abs targeting the same GPCR are grouped. Information regarding whether specific GPCR-RAMP interactions are reported in the literature is overlaid on respective cells as a solid black outline (interaction expected), a thin dashed outline (interaction not expected) or a thick dashed outline (interaction uncertain due to conflicting reports). White boxes indicate GPCR-RAMP pairs not tested. Green, complex detected; orange, complex not detected.

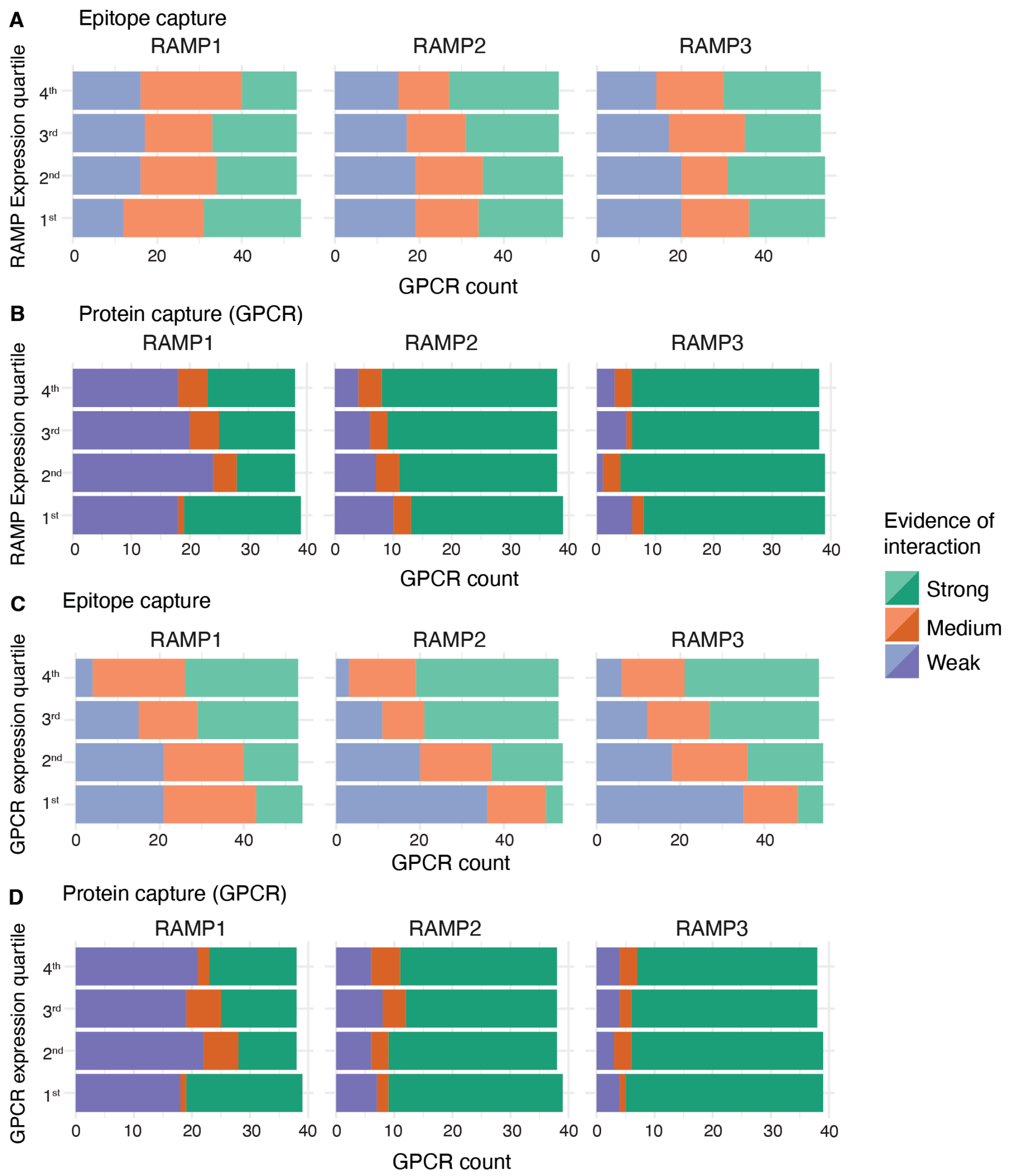


**Fig. S6. Analysis of expression level bias for detecting GPCR-RAMP interactions.**

Stacked horizontal bar plots demonstrate the distribution of GPCR-RAMP interactions by evidence class (green, strong; orange, medium; purple, weak). Strong: >66% passing capture-detection schemes. Medium: 33-66% passing capture-detection schemes. Weak: <33% passing capture-detection schemes. The plots are segmented based on RAMP expression levels (**A, B**) or GPCR expression levels (**C, D**). GPCR-RAMP complexes were identified with epitope-based capture of the GPCR and RAMP (**A, C**) or protein-specific capture of the GPCR (**B, D**). There were five unique epitope-based capture-detection schemes and up to six unique protein-based capture-detection schemes for each GPCR-RAMP pair studied. Protein capture data were generated only for the 154 GPCRs (out of 215) with at least one validated anti-GPCR Ab available. The highest GPCR or RAMP expression levels correspond to the fourth quartile, while the lowest correspond to the first quartile.

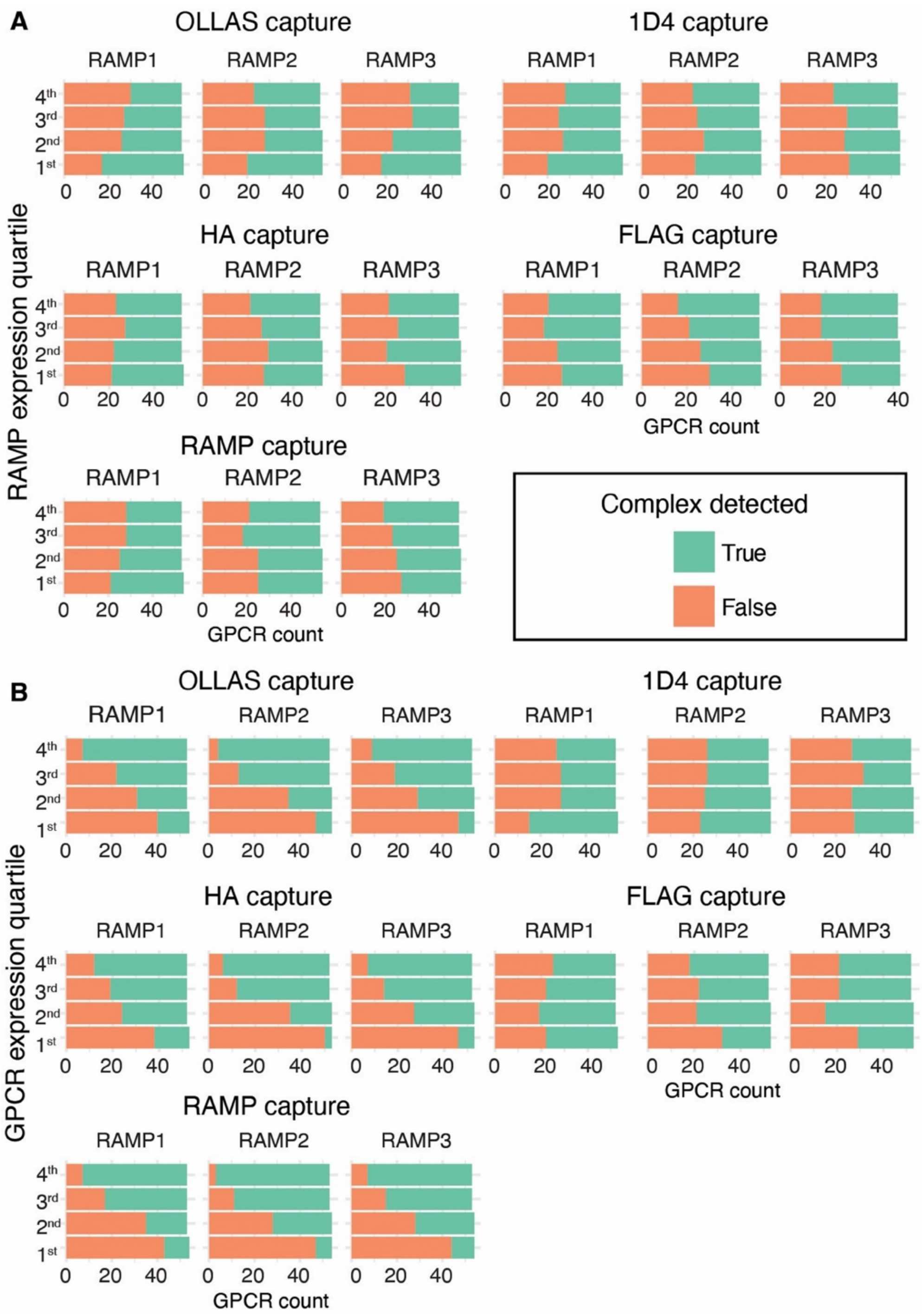


**Fig. S7. Analysis of expression level bias for epitope-based GPCR-RAMP complex detection.**

Stacked horizontal bar plots demonstrate the distribution of the total number of GPCR-RAMP complexes detected (green) or not (orange) for each RAMP with five different epitope-based capture detection schemes. The plots are segmented based on RAMP expression levels (**A**) or GPCR expression levels (**B**). *Left column*, the three capture-detection schemes are used to capture the RAMP and detect the GPCR. *Right column*, the two schemes used to capture the GPCR and detect the RAMP. The highest GPCR or RAMP expression levels correspond to the fourth quartile, while the lowest correspond to the first quartile.

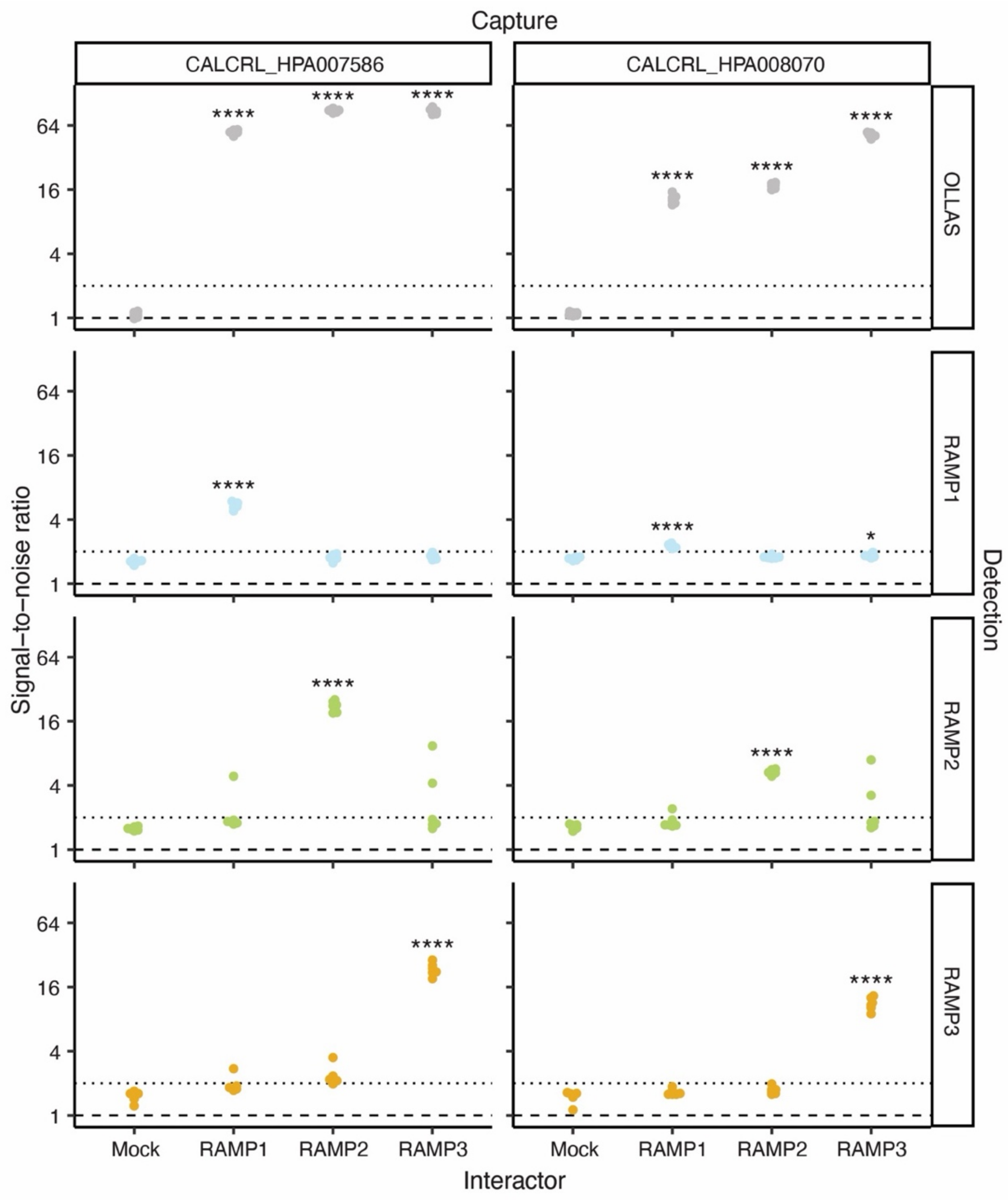


**Fig. S8. Validation of anti-RAMP Abs for detecting CALCRL-RAMP complexes.**

Validation of the detection of CALCRL co-expressed pairwise with RAMP1/2/3. Samples of solubilized membranes from Expi293F cells transfected with epitope-tagged CALCRL alone or co-transfected with each epitope-tagged RAMP were incubated with the SBA. CALCRL-RAMP complexes were captured on the beads with two previously validated anti-CALCRL HPA Abs. The complexes were detected using PE-conjugated anti-OLLAS mAb (top row) or PE-conjugated anti-RAMP1, anti-RAMP2 or anti-RAMP3 pAb (remaining rows). Statistical significance was determined by ordinary one-way ANOVA followed by Dunnett's multiple comparisons test to mock (*i.e.*, CALCRL expressed alone) (**** $p < 0.0001$, * $p < 0.01$, if not marked $p >= 0.05$). Sample names on the x-axis denote transfected RAMP name (if any). Data plotted as log2 scale. Signal-to-noise ratios: dashed line, 1; dotted line, 2. Sample sizes and p-values are listed in **Table S3**.

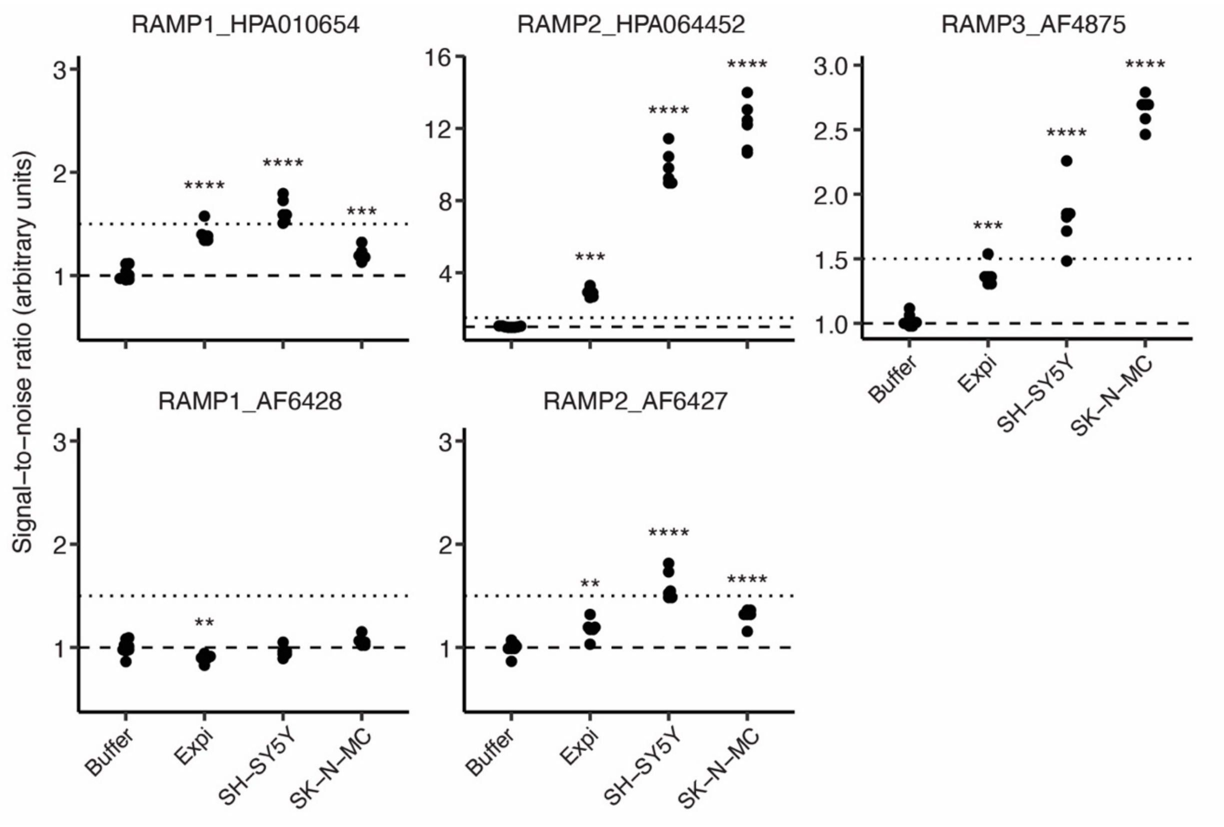


**Fig. S9. Detection of native RAMP expression in three cell lines.**

Samples of solubilized lysates from Expi293F (Expi), SH-SY5Y and SK-N-MC cells were incubated with the SBA and native RAMP1, RAMP2 and RAMP3 were captured with anti-RAMP specific Abs. The proteins were detected by PE-conjugated anti-RAMP1, anti-RAMP2 or anti-RAMP3 pAbs. Significance was determined by an ordinary one-way ANOVA with $p < 0.05$ followed by Dunnett's multiple comparison test to buffer (** $p < 0.01$, *** $p < 0.001$, **** $p < 0.0001$, $p >= 0.05$ if not marked). Data plotted as log2 scale. Signal-to-noise ratios: dashed line, 1; dotted line, 2. Sample sizes and p-values are listed in **Table S3**. Data are plotted on a log2 scale of SNR.

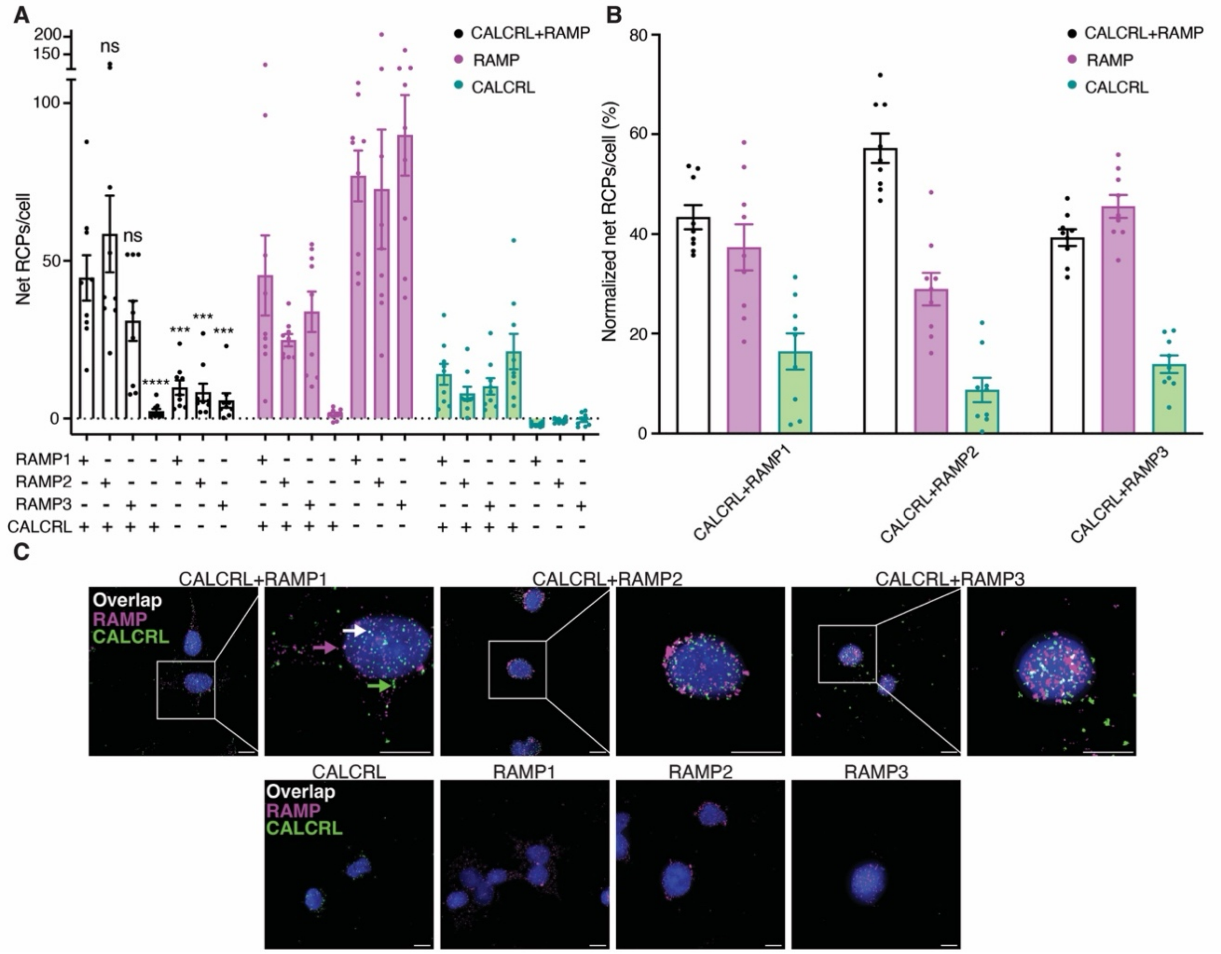


**Fig. S10. MolBoolean assay to detect CALCRL-RAMP interactions in cell membranes.**

SK-N-MC cells were incubated with anti-CALCRL primary Ab and either anti-RAMP1, anti-RAMP2 or anti-RAMP3 primary Ab and subjected to MolBoolean assay. (**A**) Signals for complexed and un-complexed CALCRL and RAMP were plotted and compared between cells stained with two primary Abs or just one. Results are shown as the number of net rolling circle amplification products (RCPs) per cell. Statistical significance was determined by one-way ANOVA followed by Dunnett's multiple comparisons test to CACLRL-RAMP1 for 'CALCRL+RAMP' (Black bars and dots) only (****$p < 0.0001$; ***$p < 0.001$; ns, not significant). Sample sizes and p-values are listed in **Table S3**. (**B**) Quantification of the percentage of RCPs out of the total number of RCPs/cell (normalized net RCPs/cell) for cells that received anti-CALCR and anti-RAMP primary Abs. (**C**) Representative Z-stack maximum projection images. Scale bars, 10 μm. Blue, DAPI; green, CALCRL puncta; magenta, RAMP puncta; white, CALCR-RAMP complex puncta. Colored arrows point to examples of puncta. RCPs from negative control MolBoolean-stained cells not incubated with any primary Ab were subtracted to obtain net RCP values. Labels on the x-axis in (**A**) and (**B**) and on top of the images in (**C**) indicate the target(s) of the included Abs. Data are from three biological replicates performed with at least three technical replicates. **Table S1.**

Antibodies and GPCRs. List of all used antibodies (Ab), their target GPCR or RAMP, and information about each GPCR. Information about which assay each Ab was used in is also specified.

(provided as a separate Excel file)

**Table S2.**

Fitting parameters of IP1 Dose-response curves. Values provided correspond to the dose-response curves presented in **Fig. S2**, computed in Prism 10. Curves were fit to a log[agonist] versus response three parameter non-linear model with least squares fit at an alpha of 0.05. Df (degrees of freedom) was computed from the number of samples and groups and indicates the number of independent pieces of information. All other values presented are the calculated best-fit values. $EC_{50}$, concentration of agonist that provokes a response halfway between the basal and maximum level of stimulation.

| | **Basal** | **$E_{max}$** | **$logEC_{50}$** | **$EC_{50}$** | **span** | **Df** |
|---|---|---|---|---|---|---|
| **HA-CALCRL-1D4 + RAMP1 (Dose-response CGRP)** | | | | | | |
| **FLAG-RAMP1-OLLAS** | 4.908 | 75.32 | -8.845 | 1.429e-9 | 70.42 | 21 |
| **3xHA-RAMP1-OLLAS** | 6.760 | 76.02 | -8.834 | 1.467e-9 | 69.26 | 21 |
| **HA-CALCRL-1D4 + RAMP1 (Dose-response Adrenomedullin)** | | | | | | |
| **FLAG-RAMP1-OLLAS** | 5.971 | 83.69 | -7.209 | 6.186e-8 | 77.72 | 21 |
| **3xHA-RAMP1-OLLAS** | 6.818 | 82.96 | -6.891 | 1.286e-7 | 76.10 | 21 |
| **HA-CALCRL-1D4 + RAMP2 (Dose-response Adrenomedullin)** | | | | | | |
| **FLAG-RAMP2-OLLAS** | 4.598 | 53.48 | -8.252 | 5.592e-9 | 48.89 | 21 |
| **3xHA-RAMP2-OLLAS** | 4.138 | 39.29 | -8.178 | 6.631e-9 | 35.15 | 21 |
| **HA-CALCRL-1D4 + RAMP3 (Dose-response Adrenomedullin)** | | | | | | |
| **FLAG-RAMP3-OLLAS** | 4.987 | 96.09 | -8.886 | 1.300e-9 | 91.10 | 21 |
| **3xHA-RAMP3-OLLAS** | 4.634 | 94.19 | -9.036 | 9.194e-10 | 89.56 | 21 |
| **FLAG-CALCRL-1D4 + RAMP1 (Dose-response CGRP)** | | | | | | |
| **FLAG-RAMP1-OLLAS** | 7.213 | 66.32 | -9.254 | 5.570e-10 | 59.10 | 21 |
| **3xHA-RAMP1-OLLAS** | 5.488 | 61.50 | -9.304 | 4.966e-10 | 56.01 | 21 |
| **FLAG-CALCRL-1D4 + RAMP1 (Dose-response Adrenomedullin)** | | | | | | |
| **FLAG-RAMP1-OLLAS** | 7.310 | 67.88 | -7.646 | 2.257e-8 | 60.56 | 21 |
| **3xHA-RAMP1-OLLAS** | 6.124 | 62.39 | -7.461 | 3.456e-8 | 56.27 | 21 |
| **FLAG-CALCRL-1D4 + RAMP2 (Dose-response Adrenomedullin)** | | | | | | |
| **FLAG-RAMP2-OLLAS** | 4.698 | 91.63 | -8.271 | 5.359e-9 | 86.93 | 21 |
| **3xHA-RAMP2-OLLAS** | 4.202 | 79.70 | -8.266 | 5.417e-9 | 75.70 | 21 |
| **FLAG-CALCRL-1D4 + RAMP3 (Dose-response Adrenomedullin)** | | | | | | |
| **FLAG-RAMP3-OLLAS** | 6.015 | 95.66 | -9.051 | 8.900e-10 | 89.64 | 21 |
| **3xHA-RAMP3-OLLAS** | 3.705 | 104.3 | -9.075 | 8.412e-10 | 100.6 | 21 |

**Table S3.**

Statistical test parameters, sample sizes, and p-values. The figure panel and test type are specified. ANOVA test results are from the aov() and summary() functions. The tables contain the capture method (Capture), degrees of freedom (Df), sums (Sum Sq) and means of squares (Mean Sq), the test statistic (F value), and p-value (Pr(>F)). Dunnett test results are from the DunnettTest() function, testing against samples with mock transfection or empty samples. The table contains the sample type being tested against (interactor or gpcr_interactor), the capture method (capture), detection method (detection), the number of samples for each sample type being tested (n), the difference in observed means (diff), the 95% confidence interval (lwr.ci, upr.ci) and the p-value (pval). Wilcoxon rank-sum test results are obtained from the wilcox.test() function. The table contains the GPCR being tested (gpcr), the capture method (capture), the p-value (p.value) and the test statistic (statistic). All statistical analysis was carried out in R.

(provided as a separate Excel file)

**Table S4.**

Thresholds for GPCR-RAMP complex detection for each epitope-based capture-detection scheme, per RAMP. Thresholds were determined by the intersection of specificity and sensitivity curves, constructed based on the GPCR-RAMP interactions reported in the literature, and are reported as the R.Z-score. Sensitivity and Specificity values at the selected threshold are reported as a probability between 0 and 1.

| Interaction | Capture | Detection | Threshold | Sensitivity | Specificity |
|---|---|---|---|---|---|
| **RAMP1** | 1D4 | OLLAS | -0.60 | 0.55 | 0.62 |
| **RAMP2** | 1D4 | OLLAS | 0.18 | 0.62 | 0.64 |
| **RAMP3** | 1D4 | OLLAS | 0.92 | 0.56 | 0.50 |
| **RAMP1** | FLAG | OLLAS | -0.65 | 0.64 | 0.69 |
| **RAMP2** | FLAG | OLLAS | 0.11 | 0.57 | 0.55 |
| **RAMP3** | FLAG | OLLAS | 0.40 | 0.78 | 0.75 |
| **RAMP1** | OLLAS | 1D4 | 0.74 | 0.55 | 0.62 |
| **RAMP2** | OLLAS | 1D4 | 0.17 | 0.52 | 0.55 |
| **RAMP3** | OLLAS | 1D4 | 0.34 | 0.61 | 0.75 |
| **RAMP1** | HA | 1D4 | -0.34 | 0.36 | 0.46 |
| **RAMP2** | HA | 1D4 | 1.18 | 0.48 | 0.55 |
| **RAMP3** | HA | 1D4 | 0.67 | 0.56 | 0.25 |
| **RAMP1** | RAMP1 | 1D4 | 0.68 | 0.36 | 0.62 |
| **RAMP2** | RAMP2 | 1D4 | 3.63 | 0.43 | 0.45 |
| **RAMP3** | RAMP3 | 1D4 | 3.14 | 0.56 | 0.75 |

**Table S5.**
GPCR-RAMP complex detection outcomes for previously reported interactions. The number of GPCR-RAMP complexes that are true positives (TP), true negatives (TN), false positives (FP) and false negatives (FN) per capture-detection scheme, per RAMP based on the literature. Anti-1D4, anti-FLAG and anti-GPCR capture (GPCR capture) corresponds to anti-OLLAS detection (RAMP detection). Anti-HA, anti-OLLAS and anti-RAMP capture (RAMP capture) corresponds to anti-1D4 detection (GPCR detection). A dash indicates a threshold was not computed for the capture-detection scheme and RAMP indicated.

| Capture | | RAMP1 | RAMP2 | RAMP3 |
|---|---|---|---|---|
| **1D4** | TP | 6 | 13 | 10 |
| | TN | 8 | 7 | 2 |
| | FP | 5 | 4 | 2 |
| | FN | 5 | 8 | 8 |
| **FLAG** | TP | 7 | 12 | 14 |
| | TN | 9 | 6 | 3 |
| | FP | 4 | 5 | 1 |
| | FN | 4 | 9 | 4 |
| **HA** | TP | 4 | 10 | 10 |
| | TN | 6 | 6 | 1 |
| | FP | 7 | 5 | 3 |
| | FN | 7 | 11 | 8 |
| **OLLAS** | TP | 6 | 11 | 11 |
| | TN | 8 | 6 | 3 |
| | FP | 5 | 5 | 1 |
| | FN | 5 | 10 | 7 |
| **RAMP1** | TP | 4 | - | - |
| | TN | 8 | - | - |
| | FP | 5 | - | - |
| | FN | 7 | - | - |
| **RAMP2** | TP | - | 9 | - |
| | TN | - | 5 | - |
| | FP | - | 6 | - |
| | FN | - | 12 | - |
| **RAMP3** | TP | - | - | 10 |
| | TN | - | - | 3 |
| | FP | - | - | 1 |
| | FN | - | - | 8 |
| **anti-GPCR Abs** | TP | 3 | 13 | 15 |
| | TN | 17 | 7 | 0 |
| | FP | 2 | 6 | 5 |
| | FN | 4 | 5 | 1 |

**Table S6.**

Thresholds for GPCR-RAMP detection for each protein-based capture scheme. The table lists all anti-GPCR Abs used, sorted by GPCR name alphabetically. The threshold and interaction results for each RAMP are provided.

(provided as a separate Excel file)

**Table S7.**

Summary of GPCR-RAMP complexes detected for each GPCR with each RAMP. The first sheet lists the evidence for interaction (strong, medium, weak) for each GPCR-RAMP pair tested by epitope-based capture (215 GPCRs) and protein-based capture (154 GPCRs). Information on whether a particular interaction is expected based on the literature is also included. Strong evidence corresponds to >66% passing capture-detection schemes. Medium evidence corresponds to 33-66% passing capture-detection schemes. Weak evidence corresponds to <33% passing capture-detection schemes. The second sheet lists the number of passing capture-detection schemes for each GPCR-RAMP pair tested. The third sheet provides the number of passing capture-detection schemes for each GPCR-RAMP pair tested as a fraction out of the total number of capture-detection schemes for the approach specified. There are five capture-detection schemes in total for all epitope-based values. The number of total capture-detection schemes for protein-based capture varies from zero to six.

(provided as a separate Excel file)

**Table S8.**

Native GPCR-RAMP complexes detected by SBA assay in three human cell lines. Complexes were captured with anti-GPCR antibodies from the Human Protein Atlas and detected with an anti-RAMP antibody specific to each RAMP. Value are Robust Z-scores, calculated separately for each detection method.

| **RAMP1 detection** | | | | |
|---|---|---|---|---|
| **GPCR** | **Antibody** | **Expi** | **SH-SY5Y** | **SK-N-MC** |
| **ADGRF5** | HPA065251 | 6.89 | 25.16 | 26.59 |
| **CCKAR** | HPA073900 | 15.60 | 10.72 | 7.67 |
| **CCKAR** | HPA051284 | | 4.58 | 4.61 |
| **CHRM4** | HPA072083 | 16.32 | 13.18 | 4.06 |
| **DRD5** | HPA065783 | 3.88 | | |
| **GLP1R** | HPA065175 | | 4.03 | 4.57 |
| **GLP2R** | HPA027929 | | 5.83 | |
| **GPR20** | HPA059117 | 5.73 | | |
| **GPR37** | HPA068009 | | 3.55 | |
| **HRH4** | HPA035009 | 3.54 | 5.09 | |
| **PTGDR** | HPA049668 | 3.91 | 4.99 | 3.71 |
| **VIPR2** | HPA062707 | 4.25 | | |

| **RAMP2 detection** | | | | |
|---|---|---|---|---|
| **GPCR** | **Antibody** | **Expi** | **SH-SY5Y** | **SK-N-MC** |
| **ADGRF5** | HPA065251 | | 10.60 | 6.41 |
| **CCKAR** | HPA073900 | 14.74 | 8.99 | 6.08 |
| **CHRM4** | HPA072083 | 10.80 | 8.92 | |
| **CRHR2** | HPA073345 | 13.50 | | |
| **DRD5** | HPA065783 | 7.68 | 5.74 | 5.74 |
| **DRD5** | HPA013797 | 12.20 | | 7.34 |
| **GCGR** | HPA077370 | | | 3.54 |
| **GLP1R** | HPA065175 | | 4.14 | 3.97 |
| **GLP2R** | HPA027929 | | 4.50 | |
| **GPR37** | HPA068009 | 3.84 | | |
| **HRH4** | HPA035009 | | 4.59 | |
| **PTGDR** | HPA049668 | 3.81 | 5.75 | |
| **SCTR** | HPA007312 | 4.92 | | |
| **VIPR2** | HPA062707 | 4.57 | | |

| RAMP3 detection | | | | |
|---|---|---|---|---|
| **GPCR** | **Antibody** | **Expi** | **SH-SY5Y** | **SK-N-MC** |
| **ADGRF5** | HPA065251 | 4.04 | 8.72 | 6.39 |
| **ADRA2C** | HPA057688 | | | 3.91 |
| **ADRB3** | HPA065408 | | 4.80 | 9.56 |
| **CCKAR** | HPA073900 | 11.69 | 6.94 | 4.83 |
| **CCKAR** | HPA051284 | | 3.79 | |
| **CHRM4** | HPA072083 | 9.11 | 7.23 | |
| **CXCR2** | HPA031999 | | | 4.02 |
| **DRD5** | HPA065783 | 3.64 | | |
| **GLP1R** | HPA065175 | | 4.09 | 4.20 |
| **GLP2R** | HPA027929 | | 5.45 | |
| **GPR37** | HPA068009 | 3.58 | | |
| **GPR4** | HPA019207 | | | 8.56 |
| **GPR61** | HPA026088 | 4.06 | | 7.27 |
| **GRM2** | HPA027868 | 4.00 | 4.59 | 4.14 |
| **HCAR2;HCAR3** | HPA028660 | | 3.99 | |
| **HRH4** | HPA035009 | | 4.24 | |
| **HTR1F** | HPA005555 | | 4.24 | 5.85 |
| **MCHR2** | HPA030115 | 9.89 | 5.91 | |
| **OPN3** | HPA049243 | | 3.54 | 6.31 |
| **OPRM1** | HPA067435 | | 3.74 | 5.38 |
| **PTGDR** | HPA049668 | 3.72 | 4.80 | |
| **PTH2R** | HPA010534 | | | 6.28 |
| **S1PR1** | HPA048588 | | | 3.85 |
| **SCTR** | HPA007312 | 3.96 | | |
| **SSTR4** | HPA064252 | 3.98 | 4.07 | |
| **VIPR2** | HPA062707 | 4.34 | | |
